# Limb assignment, not spatial remapping, underlies tactile crossing effects

**DOI:** 10.1101/2024.02.01.578364

**Authors:** Tobias Heed, Julia Burbach, Nele Rödenbeck, Boukje Habets, Xaver Fuchs

**Author notes:** Correspondence: Tobias Heed.

## Abstract

Reduced tactile-spatial accuracy with crossed versus uncrossed limbs is considered key evidence for automatic remapping of touch location from skin-based to 3D-spatial codes. Yet, typical tasks use only few stimuli, potentially confounding categorical choice of a touched limb with precise stimulus location. Over five experiments, participants judged whether two touches occurred on the same limb (anatomical) or on the same side of space (external-spatial). Crossing consistently impaired performance, but more under external-spatial than anatomical instructions — contrary to what automatic remapping predicts. Limb assignment errors occurred equally frequent for arm–leg as arm–arm pairs, implying confusion about limb identity, not spatial position. With mouse-based, precise localization, crossing again impaired limb assignment, mainly for stimuli distal to the crossing point, whereas precision was unaffected: localization scatter did not increase, and showed no bias toward the limb’s usual side of space. Crossing effects therefore emerge from choosing a limb, not from localizing touch. We propose that they reflect limb assignment, with touch cueing the region of space it is habitually associated with, followed by selecting a limb whose habitual or current posture occupies that region, constrained by which limbs are task-relevant; precise localization is performed afterwards on the selected limb.

## Introduction

When we act in the world, our cognitive system must mediate between the three-dimensional environment and the inherently two-dimensional layout of the skin. Because our body is flexible, a skin patch’s 3D location depends on current posture. A prevalent view holds that the tactile system automatically, and by default, recodes skin-based input into an external, 3D-spatial location to efficiently enable action. Yet in many situations, the exact 3D location of a touch is less important than identifying which body part was touched – for example, a tap on the left shoulder typically prompts a leftward turn regardless of the tap’s precise position on the shoulder. Similarly, playing an instrument requires attributing tactile feedback to the correct hand and finger, often regardless of the finger’s location. For instance, a guitar player may notice that a finger is on the fret rather than beside it, or that a string was struck with too little force.

Accounts for tactile localization and body perception are often linked with the idea of a somatosensory map in primary somatosensory cortex (S1). A simple “readout” account would treat limb assignment as trivial: peripheral signals arrive in S1 in an orderly somatotopic map, so the active neural population should reveal where the touch occurred. However, multiple empirical findings challenge this simplification, showing that tactile processing engages reference frames beyond the skin and that posture can profoundly alter judgments even when the physical stimulation is unchanged (Fuchs et al., 2026).

The cutaneous rabbit illusion provides a good example for perceptual errors of continuous external localization vs. categorical limb identity. When brief stimuli are delivered at two or three distant locations, observers often perceive them as progressing along the trajectory between the outer stimuli (Geldard & Sherrick, 1972). This illusion can also occur across limbs, for instance when the stimuli are presented to an arm and a leg that are placed next to each other. Just like for stimulus sequences on a single limb, participants report stimuli in the middle of the sequence displaced towards the final stimulus (Martel et al., 2022). Importantly, observers sometimes assign the middle stimulus to the wrong limb altogether—reporting, for example, an arm stimulus as having occurred on the leg— revealing categorical errors of limb identity that cannot be explained by a simple somatotopic readout.

How may such unintuitive categorical limb errors arise? A prevalent idea has been that touch is automatically transformed into external, 3D-spatial coordinates, and that limb assignment is reconstructed from the computed stimulus location (“space-first”). For the multi-limb rabbit illusion, this implies that the stimulus is perceived spatially displaced towards a location occupied by the leg and, therefore, assigned to that leg. An alternative possibility is, however, that first the leg is chosen and the stimulus then perceived at a 3D-spatial location consistent with the chosen limb (“limb-first”).

Many experiments seem to support the first notion, that is, touch being perceived in space first and being assigned to a limb second (“space-first”). When humans judge which of two consecutive tactile stimuli, one on each hand, came first – termed a tactile temporal order judgment (TOJ) – they often indicate the wrong hand to have been stimulated first when their arms are in a crossed posture (Heed & Azañón, 2014; Shore et al., 2002; Yamamoto & Kitazawa, 2001). Several theoretical accounts for this phenomenon rely on the idea that touch location is automatically coded in 3D space. One explanation suggested that the stimulus is localized in space and then assigned to the incorrect hand due to the crossed posture (Kitazawa, 2002; Yamamoto & Kitazawa, 2001). Another explanation was that the touch’s 3D location is derived and then conflicts with the anatomical location on the skin, with the hand being placed in the space opposite of the body side to which it belongs (Badde & Heed, 2016; Shore et al., 2002). In all these theoretical accounts, errors made in choosing one of two limbs are interpreted as indicating that the skin location of tactile stimuli has been automatically and immediately recoded into a precise 3D-spatial code (Azanon et al., 2016; Heed & Azañón, 2014). Thus, a limb choice task is used to theorize about tactile localization.

Newer experiments, however, have provided evidence for the notion that limb choice precedes, rather than follows, precise spatial tactile location (“limb-first”). In one study, participants received two tactile stimuli while the two hands moved towards the torso. After the movement, they pointed to the location of the touch they believed had occurred first (Maij et al., 2020). When participants had chosen the incorrect hand, they did not point to the 3D location of the stimulus that had occurred at that hand. Rather, they pointed to where the (incorrectly chosen) hand had been at the time the (correct) first stimulus had been presented to the other (correct) hand. In other words, participants inferred the tactile stimulus’s 3D location after they had decided on which hand it had occurred, leading them to report spatial locations that did not match the location of either stimulus they had received. If the tactile stimulus’s 3D location is determined only after the hand has been chosen, then the choice of which hand received the stimulus cannot have depended on that very 3D location.

If assigning touch to a limb does not involve 3D touch location, then what factors determine the assignment? In a multi-limb TOJ experiment, stimuli could occur at the hands and feet; two of these four locations were chosen randomly in each trial. Both arms and legs were positioned either uncrossed or crossed in different phases of the experiment (Badde et al., 2019). Surprisingly, participants regularly claimed that the first stimulus had occurred on a limb that had not been touched at all during the given trial, and just like in the multi-limb rabbit illusion, participants sometimes confused hands and feet. Limb misattributions were governed by limb type, body side, and the touched limb’s canonical side of external space—the side where that limb typically resides irrespective of current posture—rather than by the touched limb’s actual external location. Moreover, varying posture, such as placing limbs closer together or farther apart, did not modulate misattributions, as long as the limbs retained their crossed or uncrossed posture. These results suggest that crossing effects arise from conflicts among categorical features pertaining to body parts and their canonical-spatial priors without requiring precise external stimulus localization.

Accordingly, two aspects of tactile processing, precise localization vs. limb assignment have been confounded in past experimental work. Moreover, the finding that human participants confuse hands and feet in tactile perception is rather counterintuitive. It implies that tactile processing can disregard where in primary somatosensory cortex a touch arrived and that other, presumably higher cognitive processes dominate where touch is felt on the body. Most paradigms still involve explicit spatial judgments or spatially coded responses (e.g., pointing, left/right button presses) that can recruit external localization. For example, in the TOJ task, the first stimulus is on either the left or right hand and requires either a left or right response; therefore, even though the TOJ task asks a non-spatial question (“which stimulus came first”), it implies spatial stimulus and response attributes. As a result, limb assignment and continuous external localization are often confounded. What remains untested is whether crossing effects persist in a non-spatial, comparison task that minimizes explicit spatial demands and avoids effector-specific responses—thereby isolating limb assignment from stimulus localization.

Here, we designed a paradigm that directly targets limb assignment while minimizing spatial demands. Participants received two tactile stimuli on one or two limbs positioned either uncrossed or crossed and simply reported whether both stimuli occurred on the same limb or not. The present stimulus comparison task avoids any spatial biases potentially induced in paradigms that require choosing one of two stimuli because here both stimuli must be evaluated, and the response does not pertain to their spatial attributes. Moreover, responses can be given verbally or with a spatially orthogonal mapping, and the task requires neither naming nor moving the limb. Thus, the paradigm tests whether crossing effects emerge during limb-based assignment processes while no spatial stimulus attributes are implicitly or explicitly implied.

## Results

### Experiment 1: Tactile limb assignment

Fig. 1 illustrates the experimental design of Experiment 1. In each trial, participants received two tactile stimuli in brief succession that were either on one or two limbs (Fig. 1, upper vs. lower row, factor: Number of Stimulated Limbs). One tactile stimulus was near the wrist; the other stimulus was near the elbow, either on the forearm or on the upper arm. Thus, the two stimuli were either both on the same or on different limb segments (Fig. 1, light vs. dark blue arrows; factor: Number of Stimulated Segments). The hands were held either uncrossed or crossed (factor: Posture). The two stimuli of a trial were separated by a stimulus onset asynchrony (SOA) of 50, 100, 300, or 800 ms (factor: SOA). The order of the two stimuli, distal (wrist) first or elbow (proximal) first, was pseudo-randomized and not coded as a factor. For each stimulus pair, participants indicated, by lifting toes or heels, whether the two stimuli had occurred on the same limb or not. We report results as Balanced Integration Scores (Liesefeld & Janczyk, 2019, 2022), a combined measure of reaction time and performance accuracy; separate reaction time and accuracy data are available in the Supplementary Information for all experiments.

**Figure 1:**
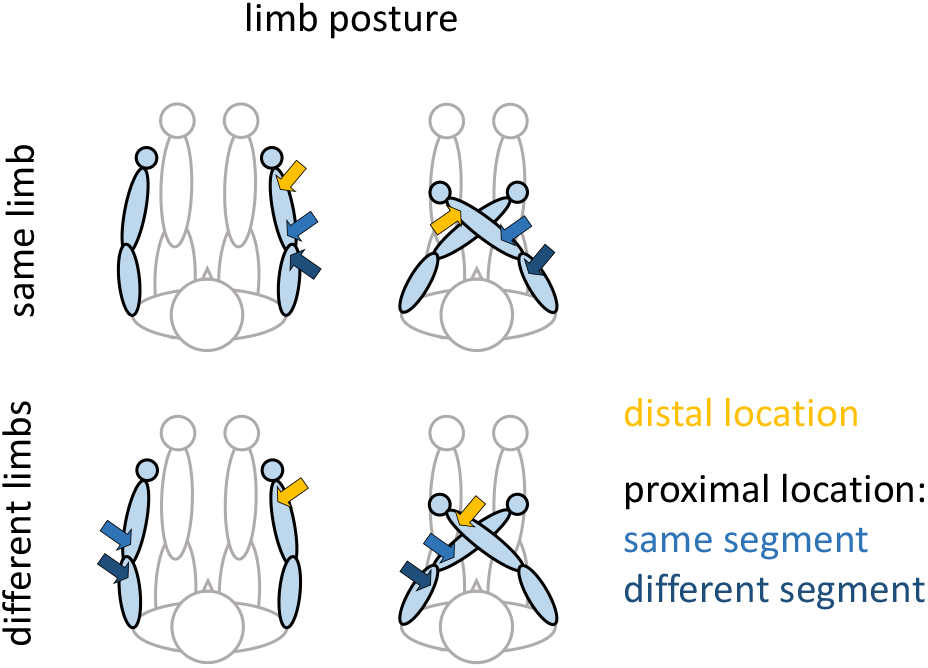
Experimental design of Experiment 1. Participants received two consecutive stimuli in each trial. Each stimulus pair comprised a distal (yellow arrows) and a proximal (light/dark blue arrows) location. Stimuli could occur on the same or on different arms, and both on the forearm (“one limb segment”, yellow plus light blue arrows) or on the fore- and upper arm (“two limb segments”, yellow plus dark blue arrows). The figure illustrates only a subset of the stimulus pairs used in the experiment to illustrate the experimental design. However, all possible stimulus combinations were realized in the experiment. For instance, the upper left panel illustrates stimulation on the right arm. Stimuli could occur first proximally and then distally, or vice versa. Moreover, all equivalent stimulus combinations on the left arm were also realized. The same rationale applies to the remaining panels.

#### Deteriorated limb assignment with crossed hands

The most notable result was a main effect of Posture, with markedly deteriorated performance when the hands were crossed rather than uncrossed (see Table 1: main effect of Posture; Fig. 2a, left vs. right panels; Fig. 2b dark vs. light blue). Thus, it was harder for participants to report whether two tactile stimuli occurred on the same limb when the limbs were crossed than when they were in a regular, uncrossed posture. Although there was large variability between participants in the raw scores (see Fig. 1b), the crossing effect was consistent across participants, evident in difference scores of uncrossed minus crossed performance >0 for all but 5 of 84 difference values in the entire sample (Fig. 2c, N=21 participants x 4 conditions; see data points below the dotted line: 2, 2, 0, 1 data points across the four depicted conditions). Post-hoc tests confirmed that the crossing effect was significant for each of the panel’s columns (all t(20) > 4.3, p < 0.001, all reported post-hoc tests FDR-corrected for multiple comparisons).

**Table 1:** ANOVA results of Experiment 1. The factor SOA has been omitted to reduce clutter. Factors: Posture (uncrossed vs. crossed); Limbs (1 vs. 2 limbs stimulated); Segments (1 vs. 2 limb segments stimulated). Results of an ANOVA including factor SOA is available in Supplementary Table 1. The effects reported here are unaffected by the additional effect and interactions with factor SOA due to the use of Type 3 sums of squares for the ANOVA. η^2^: effect size, mean Eta squared. Significant p-values are printed bold.

| Effect | df | F | $\eta^2$ | p |
| --- | --- | --- | --- | --- |
| Posture | 1, 20 | 93.91 | 0.721 | **$<0.001$** |
| Limbs | 1, 20 | 50.69 | 0.310 | **$<0.001$** |
| Segments | 1, 20 | 0.63 | 0.002 | 0.436 |
| Posture × Limbs | 1, 20 | 87.20 | 0.284 | **$<0.001$** |
| Posture × Segments | 1, 20 | 0.02 | $<0.001$ | 0.884 |
| Limbs × Segments | 1, 20 | 0.18 | 0.001 | 0.675 |
| Posture × Limbs × Segments | 1, 20 | 2.55 | 0.004 | 0.126 |

**Figure 2:**
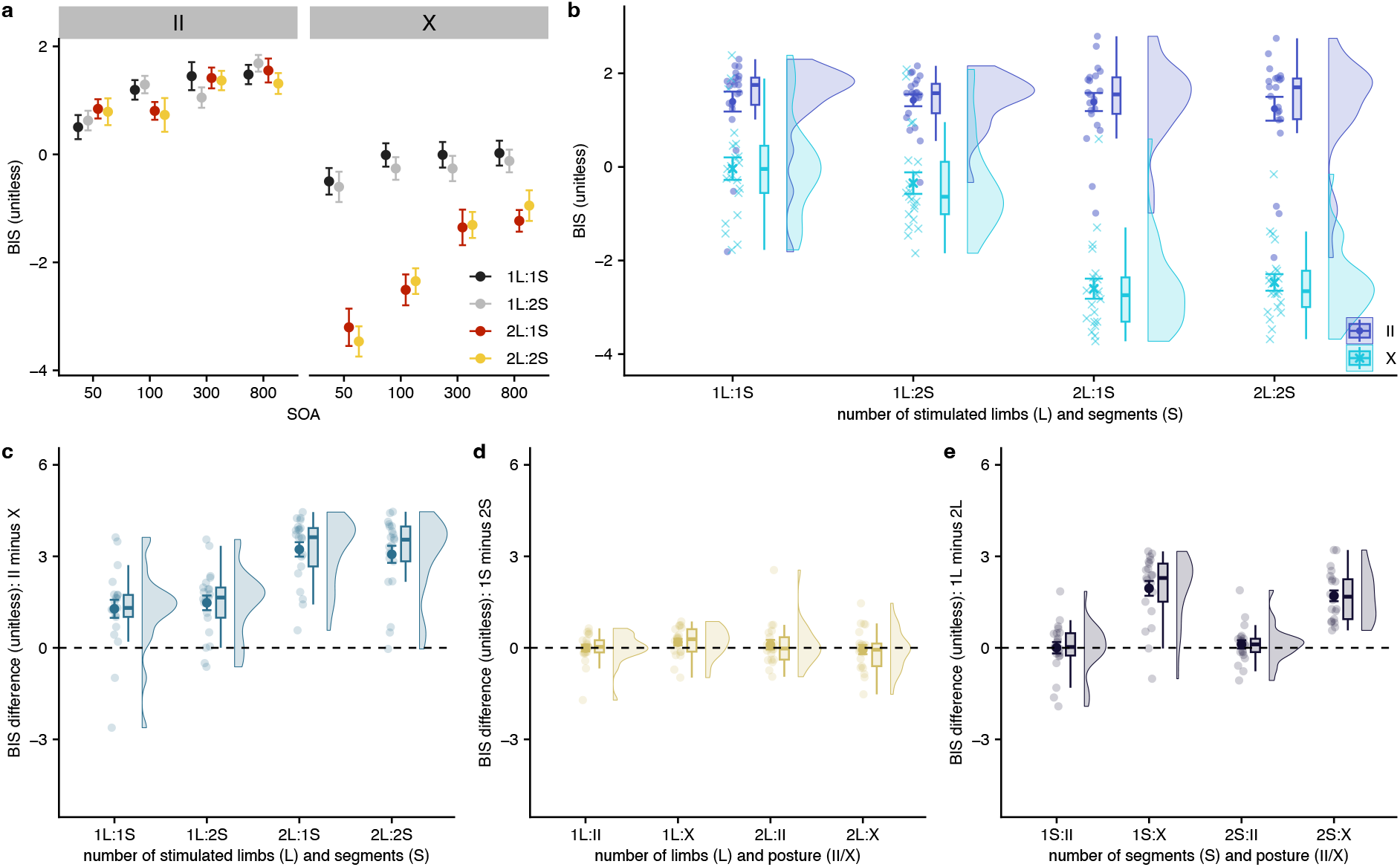
Results of Experiment 1. The Balanced Integration Score (BIS) combines reaction time and accuracy (i.e., percent correct) after by-participant normalization, thus accounting for potential trade-offs between speed and accuracy. (a) Illustration of all 4 factors of the experiment: Posture (left vs. right panel), Stimulus Onset Asynchrony (SOA), Number of Limbs and Number of Segments. Error bars denote s.d. (b) Effect of arm crossing on performance. For each condition, the left point cloud represents single participant performance, with larger/darker symbols representing the condition’s group mean and s.d. The boxplot’s upper and lower hinges indicate the 25/75th quantiles and the bold bar the condition’s median. The whiskers extend to the largest value maximally 1.5 inter-quartile ranges distant from the hinge. Outliers are not marked separately. The right violins illustrate the distribution of single participant point clouds. (c) Effect of arm posture on performance, expressed as difference scores of performance in uncrossed (II) minus crossed (X) conditions. Conventions as in *b*. (d) Effect of the number of stimulated arm segments on performance, expressed as difference scores of performance when stimuli were both presented on the forearm(s) (1S) minus on fore- and upper arm(s) (2S). Conventions as in *b*. (e) Effect of the number of stimulated limbs on performance, expressed as difference scores of performance when stimuli were both presented on the same limb (1L) minus on different limbs (2L). (c-e) Ordinate scaling is identical across panels and figures to afford visual comparisons between variables and experiments. Conventions as in *b*. II = uncrossed; X = crossed; L = limb(s); S = limb segment(s).

#### Dependence of the crossing effect on the number of stimulated limbs

Performance in the crossed posture was markedly worse when the two stimuli were presented on different limbs than on one limb; this difference was not evident in the uncrossed posture (see Table 1: interaction Posture x Limbs; Fig. 2a, black/grey vs. red/yellow data points in left vs. right panel; Fig. 2b, two leftmost [1L] vs. two rightmost [2L] conditions). Again, this effect was highly consistent across participants, with almost all difference scores of one minus two limb stimulus pairs >0 for the crossed conditions (see Fig. 2e, conditions on x-axis denoted “:X”; difference values <0: 9, 2, 8, 0 across the panel’s columns). Post-hoc tests were significant for the two crossed (both t(20) > 8, p < 0.001) but not for uncrossed conditions (both t(20) < 1, p > 0.54). The effect was evident both as longer reaction times and lower accuracy for stimuli on two limbs than one limb (see Supplementary Information, Figs. S1 and S2). It is worth pointing out that the type of error participants make is different for one- and two-limb stimulus pairs by experimental design. When participants err about one-limb stimuli, they report two stimuli on one limb as having occurred on two limbs. When they err about two-limb stimuli, they misreport two stimuli on different limbs as having occurred on one common limb. All other things equal, participants more often make the latter error, that is, they rather “perceptually merge” stimuli onto a common limb than assigning same-limb stimuli to two different body-parts. As reaction times, too, were longer for decisions about two-limb than one limb stimulus pairs to crossed limbs, the higher number of errors does not appear to be a simple bias for one over the other type of error, but more likely indicates higher difficulty of processing stimuli on two different body parts for the task of our study.

#### No effect of the number of involved limb segments

Some previous experimental results have suggested that tactile stimulus pairs are perceived as further apart when they are presented on two rather than one body part, such as on the lower arm and hand vs. on the lower arm alone (de Vignemont et al., 2009; Shen et al., 2018). Therefore, our experiment tested whether judging two stimuli as belonging to one limb is affected by the stimuli being both on the lower arm, or one on the lower and one on the upper arm. Neither the main effect of, nor any interaction with, the factor Number of Segments were significant (see Table 1, main effect of, and interactions with, Segments). Thus, whether stimuli occurred on just the forearm or across the entire arm did not significantly affect participants’ judgments. This lack of relevance can be appreciated in Fig. 2a (lighter vs. darker colors) across the full dataset, in Fig. 2b (left vs. right condition pairs) for single participants pooled across SOA, and in Fig. 2d with by-participant difference scores of the one minus two segment conditions clustering around 0; post-hoc tests for the four columns of Fig. 2d were non-significant (all t(20) < 1.8, all p > 0.36).

#### Consistent effects of SOA

Fig. 2a illustrates the results of Experiment 1 including the factor SOA. SOA affected judgments with decreasing but consistent effects of Hand Posture and Number of Limbs with increasing SOA. In the TOJ task, in which participants must decide which of two stimuli came first, many participants exhibit a judgment reversal that is maximal around an SOA of ~100 ms (Yamamoto & Kitazawa, 2001). Such a reversal was not evident in the present experimental paradigm (additionally see Supplementary Figure 2 for a summary report of accuracy alone, in which such a reversal effect is absent as well). Given the consistency of all other effects across all levels of SOA, we pooled across SOA in the previous paragraphs to reduce the number of factors in our statistical models. The full ANOVA results including factor SOA are reported in Supplementary Table 1.

### Experiment 2: Anatomical vs. external response coding

Experiment 1 demonstrated that limb crossing impairs the ability of human participants to correctly assign two tactile stimuli to the limb(s) on which they occurred. Recall that, based on the idea that the external, 3D location of a touch is automatically constructed, crossing effects have often been attributed to a conflict between anatomical/somatotopic vs. external-spatial coding (e.g., Azanon et al., 2016; Badde & Heed, 2016; Shore et al., 2002). When the hands are crossed, the anatomical location of the stimulus (e.g., right hand) conflicts with its spatial location (here, left side). In our limb identification task, crossing the hands creates situations where similar conflicts could be relevant: in the crossed posture, two stimuli on the same limb are located on different sides of space; for example, the proximal stimulus of the right arm lies on the right while the distal stimulus lies on the left side of space (see Fig. 3, right upper panel). Thus, the external locations of the two stimuli could potentially support a classification as “different limbs”, because their locations are on opposite sides of space. The underlying assumption of such a hypothesis is that the representation of tactile stimuli depends, at least in part, on the current spatial configuration of the body – an idea that has been prominent in theories of tactile-spatial processing (Badde & Heed, 2016; Kitazawa, 2002; Shore et al., 2002). Conversely, when two stimuli occur on two crossed limbs, they are located on the same side of space (see Fig. 3, right lower panel). This “same space” property of the two stimuli may support their misclassification as occurring on the same limb.

**Figure 3:**
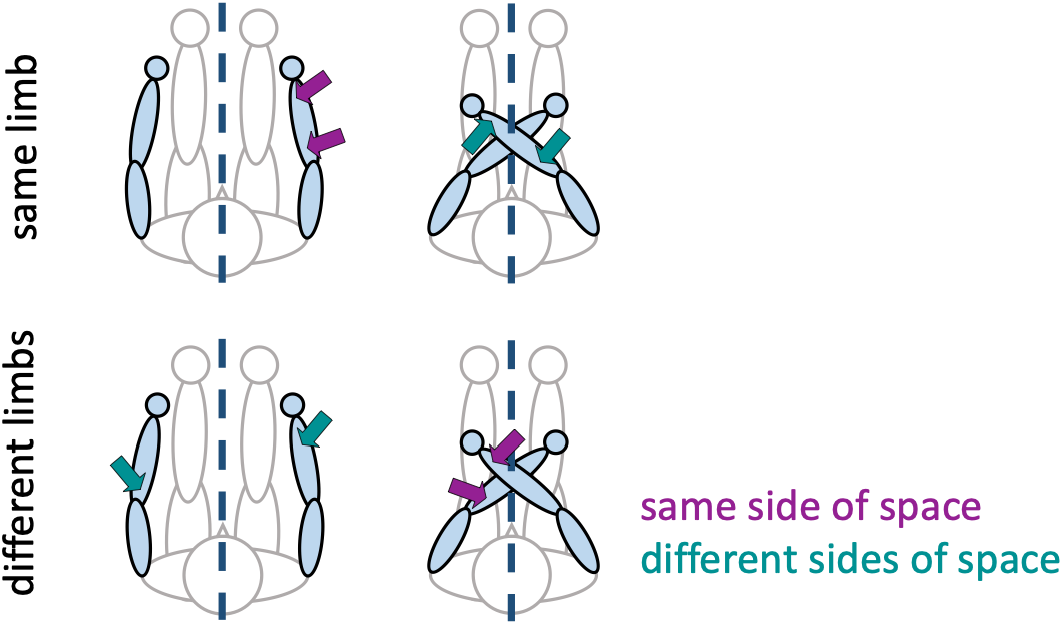
Experimental design of Exp. 2. Stimulus pairs were always presented on the forearms. Participants performed two tasks on the stimuli. In the first task, they decided whether the stimulus pair had occurred on the same limb (upper row) or not (lower row), just as in Exp. 1. In the second task, they decided whether the two stimuli had occurred on the same side of external space (purple arrows, both on one side of the dotted midline) or not (green arrows, each on a different side of the dotted midline). As for Exp. 1, we illustrate only a subset of stimulus pairs here, but all possible stimulus combinations were used in the experiment (e.g., upper left panel, stimuli on the left arm; equivalently for the remaining panels).

However, recall that recent experiments have put exactly this dependence of tactile processing on 3D space into question, suggesting instead that crossing effects result from processing of limb rather than stimulus location. The present experiment, therefore, aimed to test whether the crossing effect observed in Experiment 1 should be interpreted as indicating a modulation of limb assignment by tactile-spatial stimulus location. If the idea is correct that tactile location is derived automatically and immediately, and that the effects of limb crossing on tactile limb identification are due to anatomical vs. 3D-spatial conflict, then the crossing effect should not occur, or be significantly reduced, when participants judge stimuli regarding their spatial location. This is because the predominant, spatial code should be readily available and, thus, support fast and correct responses. In Experiment 1, participants made anatomical limb decisions (“stimuli on 1 vs 2 limbs?”). If it is the external-spatial location of the stimuli that induces the crossing effect by conflicting with these task instructions, then this conflict should be reduced when participants judge stimuli according to whether they lie on the same side of space or not. We tested this prediction in a new participant sample. Given that SOA did not modify the direction of any other experimental factor in Experiment 1, we used only a short SOA of 50 ms, because effects of Experiment 1 were strongest with short SOAs (see Fig. 1a). Moreover, we dropped the differentiation of 1 vs. 2 limb segments, because it had not shown any effect in Experiment 1. Thus, participants made either anatomical or external judgments about two tactile stimuli (factor: Instruction; levels: anatomical instruction = “were the two stimuli on the same limb or not”; spatial instruction = “were the two stimuli on the same side of space or not”); stimuli were presented either on one or on two limbs (factor: Number of Limbs) and the arms were either uncrossed or crossed (factor: Posture). The response mode was identical to that of Experiment 1, so that the spatial character of the experiment was induced solely via instruction and not via spatial response coding.

#### Larger crossing effect for external than for anatomical instruction

Whereas the account of automatic transformation of touch location into 3D space predicts a reduced crossing effect for the external instruction, the crossing effect was significantly larger when participants had to consider spatial stimulus location identity rather than anatomical limb identity (see Table 2: main effect of Instruction and interaction Instruction x Posture; Fig. 4a: condition-wise data; Fig. 4c: difference scores). As in Experiment 1, the crossing effect was present in virtually all participants and conditions, with only two of 108 difference values <0 (Fig. 4c, N=27 x 4 conditions, difference values <0: 1, 1, 0, 0 across the panel; corresponding post-hoc tests all t(26) > 7, p < 0.001). Fig. 4d illustrates that the difference between the two types of instruction was evident in the crossed, but not the uncrossed posture (difference values <0 across the panel: 17, 2, 13, 7; post-hoc tests for II, t(26) < 1.3, p > 0.28; for X, t(26) > 3.0, p < 0.009). Moreover, the Instruction effect with crossed limbs was larger when stimuli had occurred on one limb, evident in both the interaction of Instruction x Number of Limbs in the ANOVA (see Table 2) and a post-hoc test between the 1L and 2L crossed conditions (:X conditions in Fig. 4d, t(26) = 3.80, p = 0.005, corrected for all 6 possible comparisons in the panel). The larger crossing effect under external than anatomical instructions was consistent across the participant sample for stimulus pairs on one limb, with only 2 of 27 difference scores <0 (see Fig. 4d, 2^nd^ column). For stimulus pairs on two limbs, the effect was slightly less consistent, with 7 of 27 difference scores <0 (see Fig. 4d, 4^th^ column). For one-limb pairs on crossed limbs, the correct response under external instructions is that the two stimuli occurred on different sides of space. Participants, however, appear to be strongly biased to incorrectly respond that the two stimuli occurred on the same side of space. presumably because they are misled by the fact that the two stimuli were on one common limb. There are two interpretations for this finding. First, the error pattern mirrors that in the original limb assignment task in that participants overall tend to respond “same”. Thus, it may be a general phenomenon that participants perceive tactile stimuli to be similar when limb crossing makes them uncertain about the stimulus locations. Second, the error pattern indicates that participants find it hardest to respond correctly when they should reject the question (“are the two stimuli on the same side of space?”) and the anatomical location of the two stimuli implies identity (same limb). Thus, participants may be using dominant anatomical information despite external instructions. This interpretation emphasizes the relevance of anatomical information for the task and, in turn, does not fit with the idea that the external-spatial location of the two stimuli is readily available.

**Table 2:**
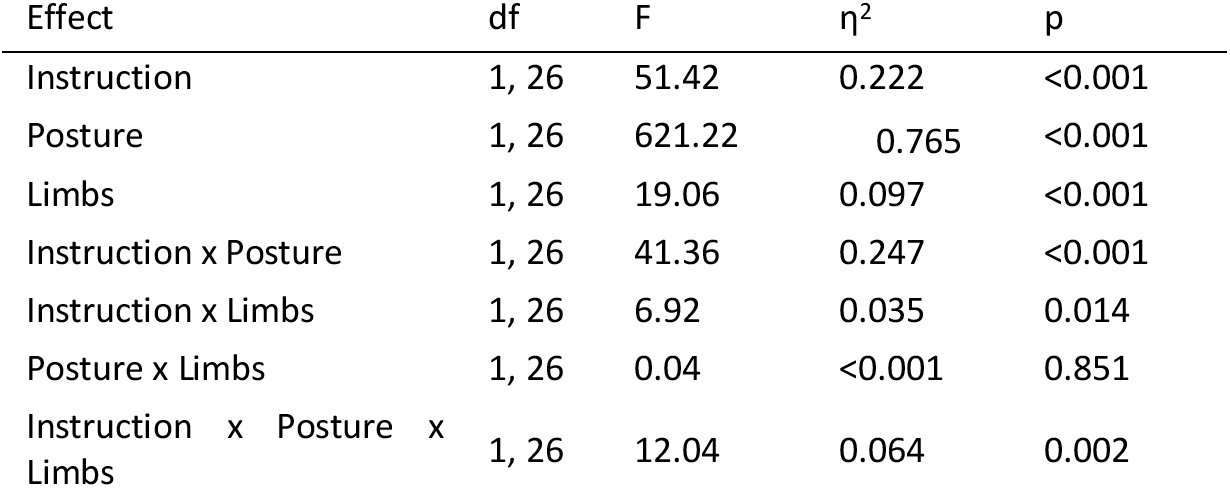
ANOVA results of Experiment 2. Factors: Instruction (anatomical vs. spatial); Posture (uncrossed vs. crossed); Limbs (1 vs. 2 limbs stimulated). Significant p-values are printed bold. See text for details.

**Figure 4:**
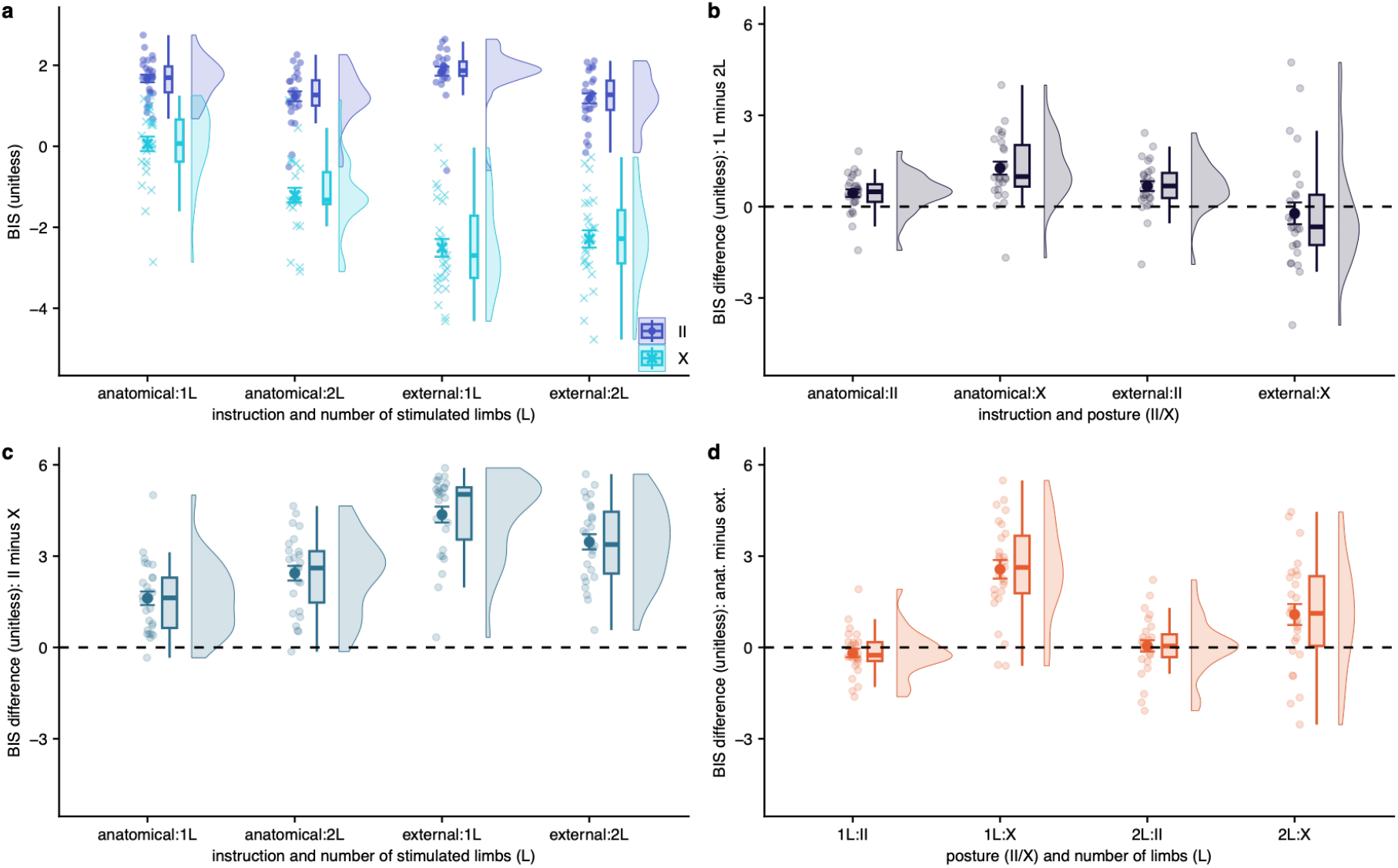
Results of Experiment 2. (a) Performance, expressed as BIS, in uncrossed and crossed postures across instructions and number of limbs. (b) Effect of the number of stimulated limbs on performance, expressed as difference scores of performance when stimuli were both presented on the same limb (1L) minus on different limbs (2L). (c) Effect of arm posture on performance, expressed as difference scores of uncrossed (II) minus crossed (X) conditions. (d) Effect of instructions on performance, expressed as difference scores of performance with anatomical (anat.) minus with external (ext.) instructions. Conventions, abbreviations as in *Fig. 2*. Coloring of analyses equivalent across experiments (panels a, b, c) match the coloring of Fig. 2.

#### Replication of the relevance of the number of stimulated limbs

Experiment 2 replicated the main effect of the number of stimulated limbs, with better performance for stimulus pairs presented to one common limb than to two different limbs, albeit only for some conditions, as evident in the interaction of Instruction x Number of Limbs and Instruction x Posture x Number of Limbs (see Table 2). This dependence on one-limb vs. two-limb stimulus pairs can be appreciated in Fig. 4c, which shows that the difference between crossing effects (II minus X performance) due to instruction is larger for one than two-limb stimuli. Fig. 4b presents the difference scores of performance for one and two-limb pairs; it is evident that the advantage of one-limb pairs is consistent for all conditions but the crossed posture under external instruction (difference values <0: 5, 1, 3, 18 of 27, across the panel’s columns; post-hoc tests for columns 1-3, t(26) > 3.5, p < 0.003; column 4, t(26) = 0.62, p = 0.54). This effect pattern appears sensible under the premise that the anatomical identity of the two stimuli (“same limb”) is consistent with the anatomically instructed task (“stimuli on same limb?”) but does not support the decision required in the externally instructed task (“stimuli on same side of space?”). Thus, it supports the conclusion that limb assignment relies on an anatomical code but is inconsistent with the traditional idea that tactile location is readily recoded into a 3D-spatial code.

### Experiment 3: Anatomical modulators and generalization to other body parts

Under the traditional hypothesis that crossing effects reflect the automatic use of external-spatial stimulus locations, external instructions should have ameliorated the crossing effect in Experiment 2. The greater crossing effect under external than anatomical instructions is, therefore, incompatible with the idea that the effect emerges because participants prioritize automatically derived external stimulus locations when they determine which limb was touched. This conclusion is in line with evidence that crossing effects in tactile TOJ may be unrelated to the external location of tactile stimuli (Badde et al., 2019; Maij et al., 2020). This, in turn, raises the question which other factors are responsible for tactile crossing effects. Experiments with the TOJ paradigm have suggested that crossing effects are due to multiple anatomy-related stimulus characteristics, for instance the type of stimulated body part, such as a hand vs. a foot, and the space related to the limb’s regular location, such as the right side of the body for the right hand (Badde et al., 2019).

Our paradigm lends itself to testing for the potential relevance of a different type of anatomical characteristic. The skin can be segmented into dermatomes, that is, regions for which axons projecting from the tactile sensors typically enter the spinal cord in the same spinal cord segment. Responses of primary somatosensory cortex differ for stimuli originating from different leg dermatomes (Dietrich et al., 2017). It is, therefore, conceivable that this anatomically guided, neural organization affects processing also in hierarchically higher areas and modulates the assignment of stimuli to limbs in our task. Here, we tested this idea by placing the two tactile stimuli of a given trial onto skin locations considered to belong to the same dermatome vs. locations considered to belong to skin areas with no dermatomal overlap between the two stimuli. The design logic is that the required classification of two stimuli as belonging to a common limb should be facilitated by stimuli being registered within a common dermatome, whereas registering two stimuli in different dermatomes should impede “same” judgments when the two stimuli belong to the same limb and facilitate “different” judgments when the two stimuli belong to different limbs.

Extent and shapes of the dermatomes vary between people, but individual dermatome shapes cannot be easily determined. Knowledge about the layout of dermatomes has mainly come from clinical studies, and multiple dermatome maps have been suggested (see Lee et al., 2008). Besides individual differences in shape, signals from a given dermatome can enter the spine via multiple roots, and there is probably some communication of neurons belonging to neighboring dermatomes (Ladak et al., 2014). In a first set of 19 participants, we chose stimulus locations based on drawings and descriptions in Keegan & Garrett (1948) without regarding potential overlap and division across multiple roots. In a second set of 19 participants, we chose stimulus locations so that the involved dermatomes would have minimal overlap and not share any spinal cord roots (Lee et al., 2008; Schünke et al., 2018b, 2018a). We chose to present stimuli to the legs because choice of stimulus locations targeting specific dermatomes was easier for the legs than the arms. As we had stimulated the arms in both Experiments 1 and 2, Experiment 3 tested the generalization of our previous experimental findings to other body parts as a side effect. Stimulus locations for Exp. 3 are illustrated in Fig. 5.

**Figure 5:**
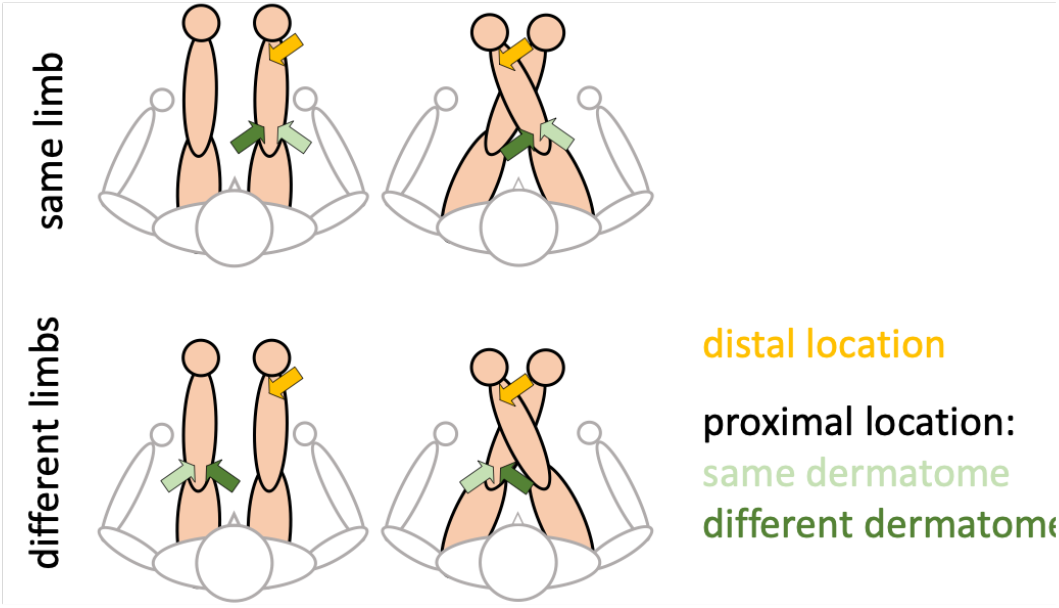
Experimental design of Exp. 3a and 3b. Stimulus pairs were presented on the legs rather than the arms. Accordingly, responses were given with the hands. There were two proximal stimulus locations; these were on the sides of the knees for Experiment 3a, and on the upper thighs for Experiment 3b. In both experiments, one proximal stimulus belonged to the same dermatome as the ankle stimulus (illustrated in light green) whereas the other belonged to a different dermatome (illustrated in dark green). The proximal stimulus locations were the only difference between Experiments 3a and 3b.

#### Generalization of the crossing effect from arms to legs

The most salient result of Experiment 3 is that the crossing effect is just as prominent for stimuli on the legs as it was for stimuli on the arm in the first two experiments (see Fig. 6a, dark vs. light blue; Fig. 6c, positive group difference of II minus X for all combinations of limbs and dermatomes, with post-hoc tests for all four conditions of Fig. 6c t(37) > 5, p < 0.001). Only few data points indicated a reversed crossing effect (Fig. 6c, data points <0, see next paragraph), suggesting that the crossing effect is a consistent phenomenon.

**Figure 6:**
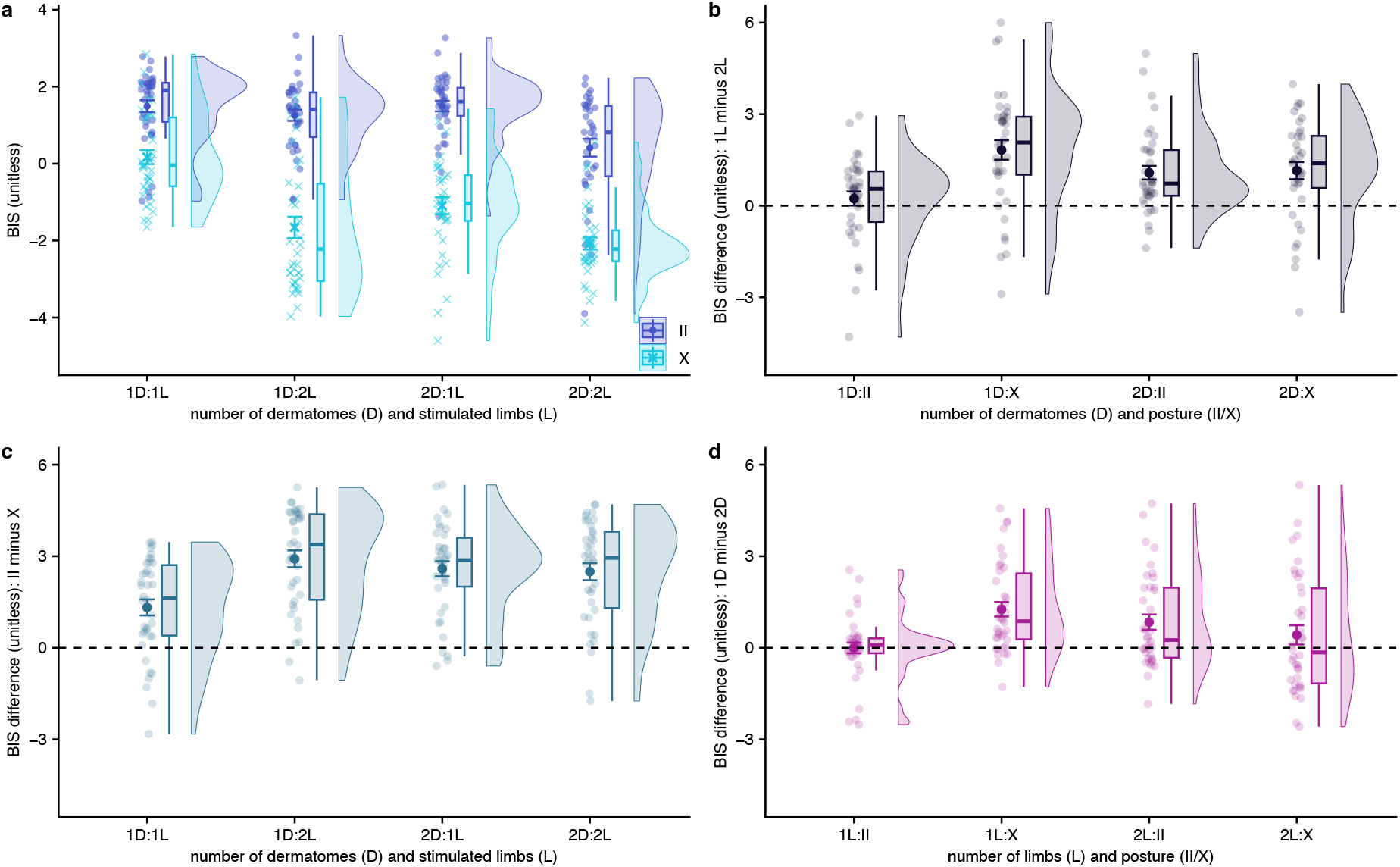
Results of Experiment 3. (a) Performance, expressed as BIS, in uncrossed and crossed postures across number of dermatomes and stimulated limbs. (b) Effect of the number of stimulated limbs on performance, expressed as difference scores of performance when stimuli were both presented on the same limb (1L) minus on different limbs (2L). (c) Effect of leg posture on performance, expressed as difference scores of uncrossed (II) minus crossed (X) conditions. (d) Effect of instructions on performance, expressed as difference scores of performance for stimulus pairs presented to one (1D) vs. two different (2D) dermatomes; note, that “1D” also refers to stimulus pairs presented to homologue dermatomes on the two body halves (1D/2L), which do not share their spinal entry point. Only stimulus pairs that share dermatome and limb (1D/1L) enter the spine at the same root. Conventions, abbreviations, and coloring of analyses equivalent across experiments (panels a, b, c) as in *previous figures*.

**Figure 7:**
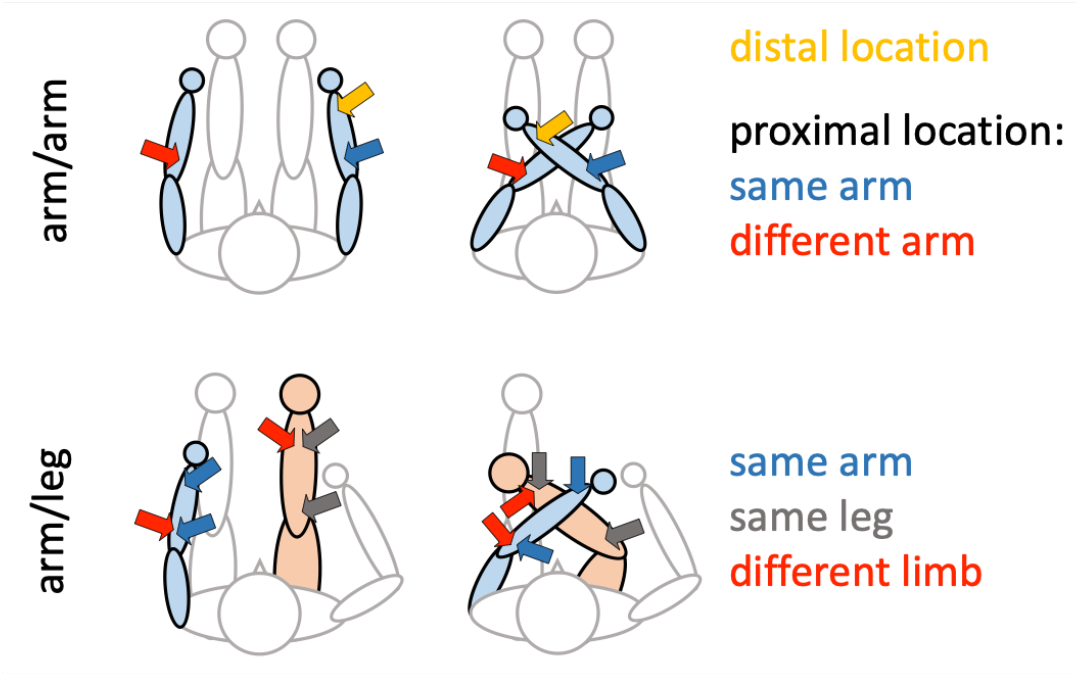
Experimental design of Exp. 4. Stimulus pairs were presented on the two arms (upper row) or on an arm and a leg (lower row). Half of participants received stimulation on the left arm/right leg, the other half vice versa. Participants gave verbal responses. For crossing arm and leg, both limbs were placed such that the end effector (hand, foot) as well as the distal stimulus were located on the side of space opposite to the limb’s body side. The red arrows in the lower row illustrate only one of two stimulus configurations for arm/leg stimulus pairs: stimuli could also occur on a distal arm and proximal leg position.

#### Reduced crossing effect for stimulus pairs within a common dermatome

The first column of figure 6c illustrates that the crossing effect was smaller for stimulus pairs presented to one dermatome. That this factor combination stands out from all others is evident in the significant 3-way interaction of Limbs x Dermatomes x Posture as well as the 2-way interaction of Limbs x Dermatomes (see Table 3). Indeed, post-hoc t-tests confirmed that the crossing effect for stimulus pairs within one dermatome of one limb differed from all other conditions (all t(37) > 4.7, p < 0.001), whereas these respective other conditions did not differ from each other (all t(37) < 1.75, p > 0.3). Moreover, 7 of the 38 participants had a reversed crossing effect, that is, a II-X difference <0, whereas this number was lower in the remaining conditions (4, 2, and 3, in the order of conditions as depicted in Fig. 6c). This dermatome effect cannot be explained by spatial distance. Both the same and different dermatome stimuli were placed far away from the respective, paired stimulus in each trial: one stimulus was always near the foot, and both the same and different dermatome stimuli were placed near the knee or on the upper thigh.

**Table 3:** ANOVA results of Experiment 3. Factors: Posture (uncrossed vs. crossed); Limbs (1 vs. 2 limbs stimulated); Dermatomes (1 vs 2 dermatomes stimulated). Significant p-values are printed bold. See text for details.

| Effect | df | F | $\eta^2$ | p |
| --- | --- | --- | --- | --- |
| Posture | 1, 37 | 125.77 | 0.494 | <0.001 |
| Limbs | 1, 37 | 32.90 | 0.161 | <0.001 |
| Dermatomes | 1, 37 | 18.73 | 0.067 | <0.001 |
| Limbs x Posture | 1, 37 | 11.50 | 0.025 | 0.002 |
| Dermatomes x Posture | 1, 37 | 4.97 | 0.008 | 0.032 |
| Dermatomes x Limbs | 1, 37 | <0.001 | <0.001 | 0.994 |
| Dermatomes x Limbs x Posture | 1, 37 | 26.37 | 0.031 | <0.001 |

Our factor Dermatome implies that stimulus pairs on two limbs can share a dermatome. However, the signals related to the two respective stimuli enter the spine on opposite body sides and, thus, do not truly share a dermatome; rather, the factor level “same” refers to dermatomes of the two body halves being homologues, which are clearly separated neurally. Only stimulus pairs that are presented to a common limb can truly share a dermatome – and it is those stimulus pairs for which limb crossing has a significantly smaller effect. Accordingly, in Fig. 6d, which displays the performance difference for stimulus pairs presented to common vs. separate dermatomes (1D vs. 2D), the difference is greatest for stimulus pairs on a common leg (1L) when the legs are crossed (Fig. 6d, 2^nd^ column, 1L:X). Still, even for one-limb, one-dermatome stimulus pairs, the crossing effect was highly significant (t(37) = 5.02, p < 0.001).

#### High variability between participants for the dermatome effect

Fig. 6d depicts the variability of the dermatome effect across participants. Comparison with the crossing effect (see Fig. 6c) illustrates that the dermatome effect is much less consistent across participants (17, 7, 15, 20 participants with values <0, in the order of columns of Fig. 6d; with N=38, the 20 participants of the 4^th^ column make up more than half of the sample; compare to values < 0 in Fig. 6c). Indeed, the 3-way interaction of Dermatomes x Limbs x Posture implies differences between the different factor combinations. Post-hoc tests revealed that the dermatome effect (1D minus 2D) was significant only for the factor combinations 1L/X (t(37) = 5.31, p < 0.001) and 2L/II (t(37) = 3.36, p = 0.004) but not the remaining combinations (p > 0.25).

The definition of dermatomes is fuzzy: For one, dermatomes differ between individuals. Moreover, input sometimes diverts to neighboring spinal cord segments, leading to overlap between neighboring dermatomes (Ladak et al., 2014; Lee et al., 2008). These factors likely affected our experiment, for instance, with placement of a stimulator at a location of dermatome overlap for some participants. The large variability and the small effect size of the dermatome manipulation may partly relate to this impossibility to determine the precise extent of the dermatomes in healthy participants with reasonable effort.

#### (Second) Replication of dependence of performance on number of stimulated limbs

As in Experiments 1 and 2, performance was slightly better, on average, when stimuli were presented on one than on two limbs. The mean size of this effect in standardized units of the BIS was similar to that of Experiment 1, but was less consistent, with more participants showing effects in the opposite direction for Experiment 3 than Experiment 1 (12, 7, 6, 9 participants, in the order of columns of Fig. 6b; post-hoc tests p = 0.35 (n.s.) for 1D:II, but p < 0.006 for all other conditions); together, these two aspects resulted in a larger effect size for the main effect of Number of Limbs in Exp. 1 than Exp. 3 (η^2^ = 0.31 vs. 0.16, see Tables 1 and 3). Still, in both Experiments 1 and 3 the Number of Limbs effect was larger for the crossed than the uncrossed condition, unlike in Experiment 2. Overall, it is evident that this effect has a smaller effect size and is less reliable than the limb crossing effect, with its effect size of η^2^ > 0.6 in all three Experiments. Experiment 1 had a higher number of trials than Experiment 2 because we pooled across 4 different SOAs; Experiment 3 had twice as many participants than Experiment 2. These differences in the amount of data may well explain the inconsistent effect in Experiment 2.

### Experiment 4: Confusion of arm and leg

We have established that limb crossing affects decisions about the identity of the stimulated limb(s) when stimuli are applied to the two arms or to the two legs during the entire experiment. Introspectively, it seems unlikely that participants would have difficulty deciding whether two stimuli occurred on a common limb if one limb is an arm and the other a leg. Arms and legs are clearly distinct regarding their position on the body as well as their functional role, and intuitively they seem easily distinguishable.

Nevertheless, previous evidence suggests the opposite: participants confused hand and foot in tactile temporal order judgments similarly as the two hands (Badde et al., 2019; Schicke & Röder, 2006) and they reported with high confidence that a single tactile stimulus presented to a hand had occurred on the foot and vice versa (Badde et al., 2019). These previous results suggest that humans can make fundamentally incorrect, but systematic, judgments about where touch occurred on the body and do not just confuse homologous limbs, such as the two arms or the two legs. Experiment 4 scrutinized this conjecture. In different phases of the experiment, we presented stimulus pairs to the two arms or to an arm and a leg. Participants responded verbally, because we wanted to avoid biasing responses towards arm or leg by using one of them to respond. While counter-intuitive, the discussed previous research led us to expect that the crossing effect observed in Experiments 1-3 should be indistinguishable for these two limb sets. We powered the experiment so that we would detect a difference between arm-arm and arm-leg performance with 90% power if one set’s crossing effect is only 70% or less as large as the other set’s. We preregistered the experiment on OSF (see Methods).

#### Generalization of the crossing effect to non-homologous limb sets

As in the previous experiments, there was a strong crossing effect (see Table 4, main effect of Posture; Fig. 8a, dark vs. light blue). In contrast, whether stimuli occurred on the two arms or on an arm and a leg did not significantly affect performance: both the main effect of Limb Set as well as all interactions with this factor were non-significant, indicating that performance was similar for the two limb sets. Fig. 8d further illustrates the difference scores between the two limb sets (arm-arm minus arm-leg) across the other experimental factors, with all distributions centering on (nearly) 0 (post-hoc tests for the panel’s four columns t(27) < 1.8, p > 0.39). Furthermore, Fig. 8c illustrates that, as in the other experiments, the crossing effect was again consistent across participants (N=28 x 4 conditions = 112 difference values; values <0: 5, 3, 2, 2 across the panel’s columns; all post-hoc tests t(27) > 4.6, p < 0.001). The result pattern of Experiment 4 is therefore as we had hypothesized and preregistered.

**Table 4:** ANOVA results of Experiment 4. Factors: Posture (uncrossed vs. crossed); Limb Set (two arms vs. an arm and a leg); Limbs (1 vs. 2 limbs stimulated). Significant p-values are printed bold.

| Effect | df | F | $\eta^2$ | p |
| --- | --- | --- | --- | --- |
| Posture | 1, 27 | 125.56 | 0.512 | <0.001 |
| Limb Set | 1, 27 | 0.79 | 0.006 | 0.381 |
| Limbs | 1, 27 | 32.31 | 0.124 | <0.001 |
| Posture x Limb Set | 1, 27 | 0.55 | 0.003 | 0.464 |
| Posture x Limbs | 1, 27 | 11.54 | 0.035 | 0.002 |
| Limb Set x Limbs | 1, 27 | 1.74 | 0.009 | 0.199 |
| Posture x Limb Set x Limbs | 1, 27 | 2.97 | 0.007 | 0.096 |

**Figure 8:**
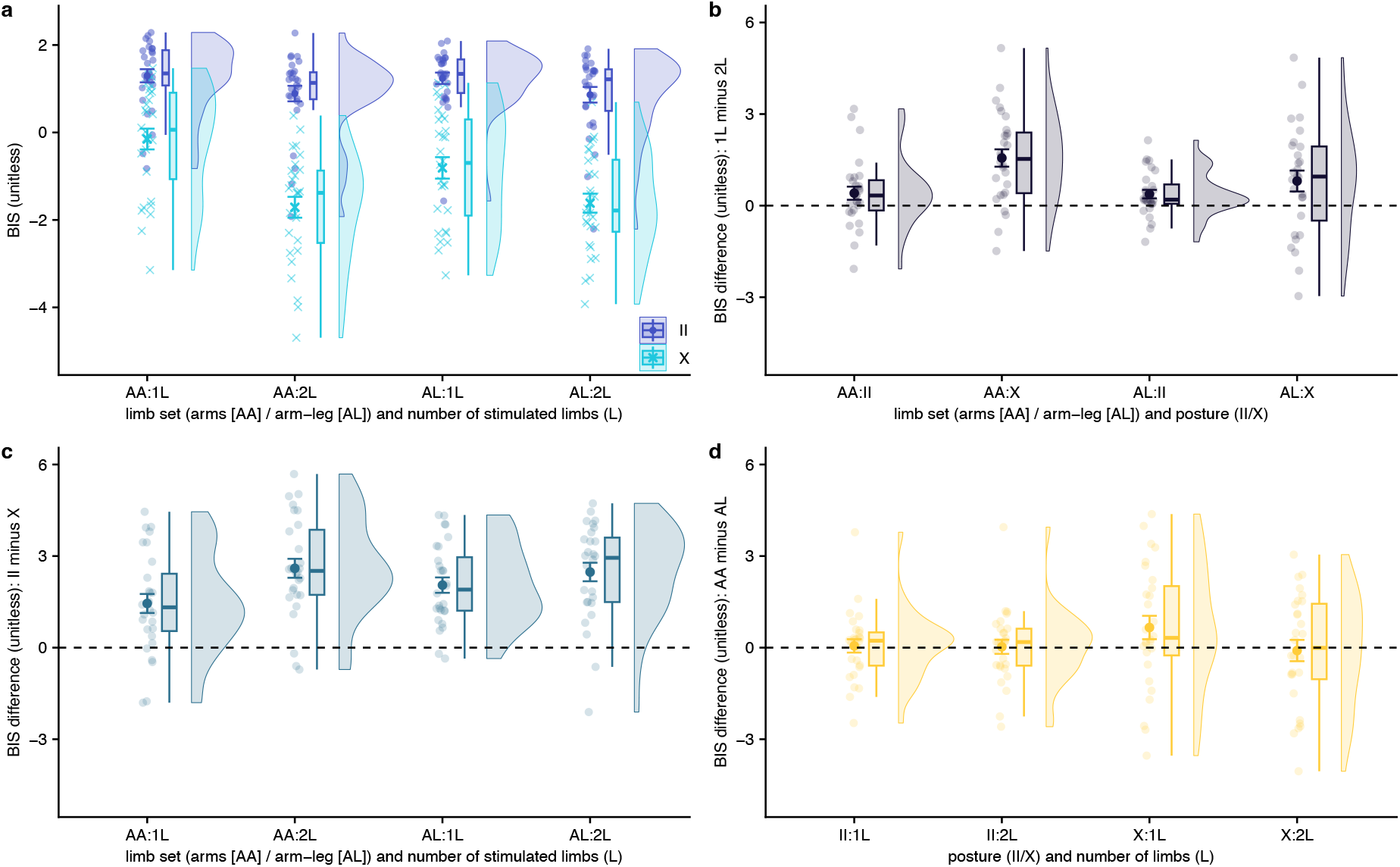
Results of Experiment 4. (a) Performance, expressed as BIS, in uncrossed and crossed postures across limb set (the two arms vs. an arm and a leg) and number of stimulated limbs (arm/leg). (b) Effect of the number of stimulated limbs on performance, expressed as difference scores of performance when stimuli were both presented on the same limb (1L) minus on different limbs (2L). (c) Effect of leg posture on performance, expressed as difference scores of uncrossed (II) minus crossed (X) conditions. (d) Effect of stimulated limbs on performance, expressed as difference scores of performance for stimulus pairs presented to the two arms (AA) vs. an arm and a leg (AL). Conventions, abbreviations, and coloring of analyses equivalent across experiments (panels a, b, c) as in previous figures.

#### (Third) Replication of dependence of performance on number of stimulated limbs

Again, performance with crossed limbs was better for stimulus pairs presented to a single limb than for those presented to two limbs, illustrated in Fig. 8b (post-hoc tests: AA:II, t(27) = 1.90, p = 0.07; remaining columns of the panel t(27) > 2.3, p < 0.04). This analysis did not differentiate between same-limb pairs presented on the arm vs. on the leg. When we split up the data of the arm-leg limb set according to whether the stimulus pair occurred on the arm, on the leg, or on both, the crossing effect was significantly smaller for arm stimulus pairs (i.e., 1L-arm) than for leg (1L-leg, p < 0.001) as well as arm-leg pairs (2L, p < 0.001).

#### Independence of particular response mode

Experiments 1-3, participants had responded with the limbs that did not receive stimulation, that is, with the feet for arm stimuli and with the hands for leg stimuli. Here, comparable effects were observed when participants responded verbally, suggesting that the crossing effects we report here occur independent of whether the task requires a motor response.

### Experiment 5: Categorical decisions vs. directed action

The first four experiments have provided evidence that human participants have substantial difficulty in assigning tactile stimuli to the limbs, and we have suggested that the results presented thus far oppose the prevalent idea that tactile stimuli are automatically recoded into an external-spatial code. However, several critical points remain that cast doubt on our conclusions.

First, one could argue that tactile stimuli are recoded as spatial locations after all, but that this spatial code is associated with greater uncertainty in crossed than uncrossed postures, for example because crossed postures are less frequent than uncrossed ones. This uncertainty may be expressed in higher variability of the estimated, spatial location, and/or in a bias of the spatial location towards the limb’s default side of space; in turn, such uncertainty may lead to slow and incorrect responses in our task. Our experiments so far have not directly assessed the perceived stimulus locations in space, and so cannot speak to this possibility.

Second, our limb assignment task has the disadvantage that participants would give a correct response if they incorrectly localized, or classified, both stimuli of a pair. For example, if we presented the two stimuli on the left arm and the participant incorrectly assigned both to the right arm, her “same limb” judgment would be correct but – unnoticeable for us – based on two incorrect, rather than two correct, assignments.

Experiment 5 addresses both concerns. Participants again received two concurrent tactile stimuli on the legs in each trial, one proximal and one distal. The legs were placed underneath a table. A monitor lay flat on the table, above the legs. Participants made goal-directed movements towards the two perceived stimulus locations of a trial by clicking on the respective locations with the mouse cursor. They performed the experiment in two different response modes (see Fig. 9). In the anatomical response mode, the monitor displayed unclothed, uncrossed or crossed legs, consistent with the participant’s own current leg posture, and participants clicked on the locations on the leg images at which they had perceived the two stimuli. In the spatial response mode, the monitor was empty, and participants clicked on the locations in space at which they thought they would see the stimulated locations if they were looking through the monitor onto their legs from directly above, as if the monitor was a window through the table surface.

**Figure 9:**
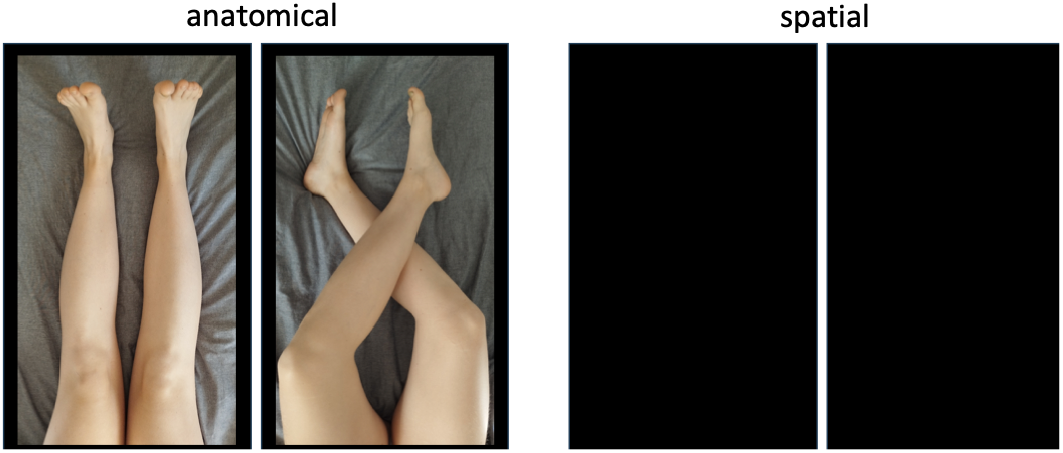
Response modes of Exp. 5. Stimulus pairs were presented on the lower legs. Participants localized the stimuli in arbitrary order by clicking with the mouse. In the anatomical response mode (left), a picture of the legs was depicted on the screen, which lay flat on top of the table under which the legs were placed. In the spatial response mode (right), participants imagined looking from straight above the empty monitor through the table surface and reported where the stimulus was located spatially if it had been visible.

#### Crossing effects with pointing responses

We partitioned each participant’s pointing responses into four clusters and classified each pointing response as an assignment to the corresponding stimulus location (see Methods). For instance, in the uncrossed leg posture, a click in the upper, left region relative to all responses of that participant was classified as a left-leg, distal assignment. This analysis strategy allowed us to infer hypothetical “same limb” vs. “different limb” responses, as they were given in Experiments 1-4. Fig. 10 depicts the results of this analysis. We present accuracy rather than BIS scores, because participants made two clicks per trial in response to two concurrent stimuli in the order of their choice, so that RT was not available for the second stimulus of a pair. A crossing effect in accuracy was evident for the anatomical and the spatial response mode, even for one-limb stimulus pairs in the former (see Fig. 10d; post-hoc tests across the four columns, p = 0.015, <0.001, <0.001, <0.001; values in opposite direction, i.e., difference < 0 across the four panels of Fig. 10d: 9, 1, 1, 1).

**Figure 10:**
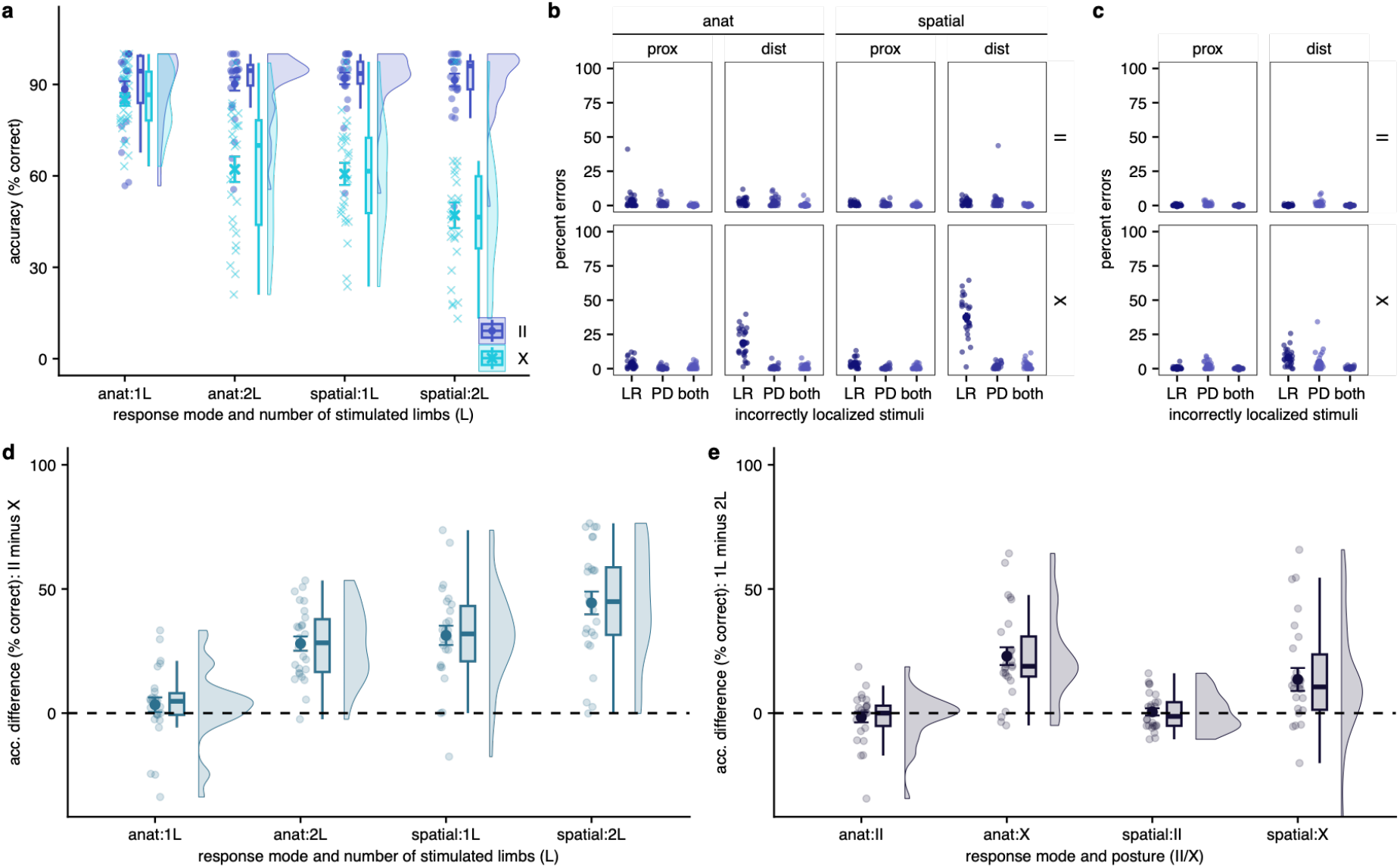
Inferred same vs different limb assignment in Experiment 5. After assigning each mouse click to one of the four possible limb locations based on the quadrant of the cursor position (see text for details), the two responses of each trial were translated into a “same” vs. “different” limb response, akin to the responses given in Experiments 1-4. (a) Percentage of correct responses in uncrossed and crossed postures across response mode (anatomical vs external) and number of stimulated limbs. (b) Error types for stimuli in the limb assignment task across response modes (anat/spatial) and stimulus region (proximal/distal). LR, response was (only) in the incorrect left/right quadarant; PD, response was (only) in the incorrect region (proximal, distal); both, response was incorrect regarding both left/right and region. (c) Error types for the single stimulus trials; these were assessed only for the spatial response mode. (d) Effect of leg posture on performance, expressed as difference scores of uncrossed (II) minus crossed (X) conditions. (e) Effect of the number of stimulated limbs on performance, expressed as difference scores of performance for stimulus pairs presented to one vs. two legs. Conventions, abbreviations, and coloring of analyses equivalent across experiments (panels a, c, d) as in previous figures.

#### Larger crossing effects for spatial than anatomical pointing responses

Moreover, the crossing effect was more pronounced for the spatial than the anatomical task (see Fig. 10c; Table 5), in line with the results of Experiment 2. One might argue that the anatomical task was easier given that depicting legs allowed precise, visually controlled pointing movements. Note, however, that the present analysis relies solely on the quadrant in which a response occurred. Thus, it refers to categorical errors in assigning a limb as the movement target, not to the spatial precision of pointing responses.

**Table 5:** GLMM results of Experiment 5. Factors: Posture (uncrossed vs. crossed); Response Mode (anatomical vs. spatial); Limbs (1 vs. 2 limbs stimulated). Significant p-values are printed bold. See text for details.

| Effect | df | $\chi^2$ | p |
| --- | --- | --- | --- |
| Posture | 1 | 42.64 | <0.001 |
| Response Mode | 1 | 5.73 | 0.017 |
| Limbs | 1 | 8.40 | 0.004 |
| Posture x Response Mode | 1 | 12.14 | <0.001 |
| Posture x Limbs | 1 | 5.91 | 0.015 |
| Response Mode x Limbs | 1 | 5.91 | 0.015 |
| Posture x Response Mode x Limbs | 1 | 1.34 | 0.246 |

#### Incorrect localization mainly for stimuli at the crossed part of the limb

When participants localized only one of the two stimuli incorrectly, the corresponding “same limb” vs. “different limbs” response is also classified as incorrect. Fig. 10b illustrates the percentage of trials in which stimuli was incorrectly localized, separately for proximal and distal stimuli. The proximal part of the lower leg, i.e. the knee, does not change its side of space in our crossed posture. The corresponding stimuli are only seldomly localized incorrectly: the error rate was 4.4% on average, and a GLMM over only proximal stimuli did not reveal a crossing effect (Fig. 10b, columns “prox”; GLMM main effect of Posture, χ^2^ = 1.54, p = 0.21; though marginally significant interaction of Posture x Response Mode, χ^2^ = 3.84, p = 0.05, with a descriptive crossing effect for the spatial but not the anatomical response mode). In contrast, for distal stimuli error rates were markedly higher in the crossed than in the uncrossed posture (Fig 10b, columns “dist”; GLMM main effect of Posture, χ^2^ = 42.95, p < 0..001). Thus, limb assignment errors in the crossed posture were particularly due to incorrect processing of the distal stimuli, which lie in the space opposite to the stimulated limb’s default space.

Fitting with the result pattern of Exp. 2, the crossing effect was larger for the spatial than the anatomical response mode (Fig. 10b, columns “dist”, lower row “X”; GLMM interaction of Response Mode x Posture, χ^2^ = 7.16, p = 0.007; mean percentage of error in the crossed posture, anatomical response mode: 20.8%; spatial response mode: 41.4%). Fig. 10b also shows how often participants confused whether a stimulus had been proximal or distal (x-axis, “PD”) and how often left/right and proximal/distal were both incorrect in a response (x-axis, “both”). It is evident that crossing affected primarily right/left decisions.

Participants also underwent some trials in which they received only a single stimulus, though only in the spatial response mode. Fig. 10c shows that the pattern of crossing effects was comparable to those evident in two-stimulus trials, with high error rates for distal, crossed stimuli. However, the overall number of errors was smaller. Thus, participants misclassify stimuli whether they localize one or two stimuli, but the two-stimulus context amplifies the effect. Thus, the limb assignment task appears to modulate but not cause the crossing effect. This finding is reminiscent of the TOJ task, which also strongly amplifies tactile judgment errors in comparison to single stimulus task settings (Badde & Heed, 2016).

#### Double errors likely reflect simple lapses

The pointing task also allowed us to estimate whether participants are likely to give responses that we interpret as correct but that derive from both stimuli having been assigned to the incorrect limb (i.e. double-incorrect resulting in a “correct” classification). We inspected how often participants had localized *both* stimuli on the wrong side in Exp. 5; this was the case in 1.3% of trials. Even though double errors were higher in the crossed posture (II: 0.8%; X: 1.7%), their absolute numbers were small compared to all other error types (see Fig. 10b) and suggests that the limb assignment task does not suffer substantially from the possibility that responses are unknowingly misclassified; as lapses are a ubiquitous phenomenon in psychophysics and psychological experimentation and are often even modeled explicitly (e.g., Wichmann & Hill, 2001).

#### Similar endpoint variability in uncrossed and crossed postures

Given that localization was incorrect mainly for distal stimuli, we analyzed the variability of the mouse cursor movement endpoints – i.e., the click locations – only for these stimuli. Endpoint variability was similar in uncrossed and crossed postures, illustrated in Fig. 11 by 95% confidence ellipses. Ellipses are remarkably similar for the two postures (comparison of s.d. in uncrossed vs. crossed postures: t(24) = 0.40, p = 0.70), suggesting that uncertainty about stimulus location cannot account for the high crossing-related error rates and RT increases of the limb assignment task. Moreover, no systematic bias towards the default limb’s space is evident. The point clouds in Fig. 11 are collapsed across left and right locations, depicted as if all stimuli had been presented on the left side of space (see Methods). A bias towards the limb’s default space would have been evident in a bulge towards the right, that is, towards the body midline, in the crossed posture. However, no such bias is evident. These results do not support the notion that the limb assignment errors observed in our task result from elevated uncertainty about the stimuli’s exact location in space in the crossed limb posture.

**Figure 11:**
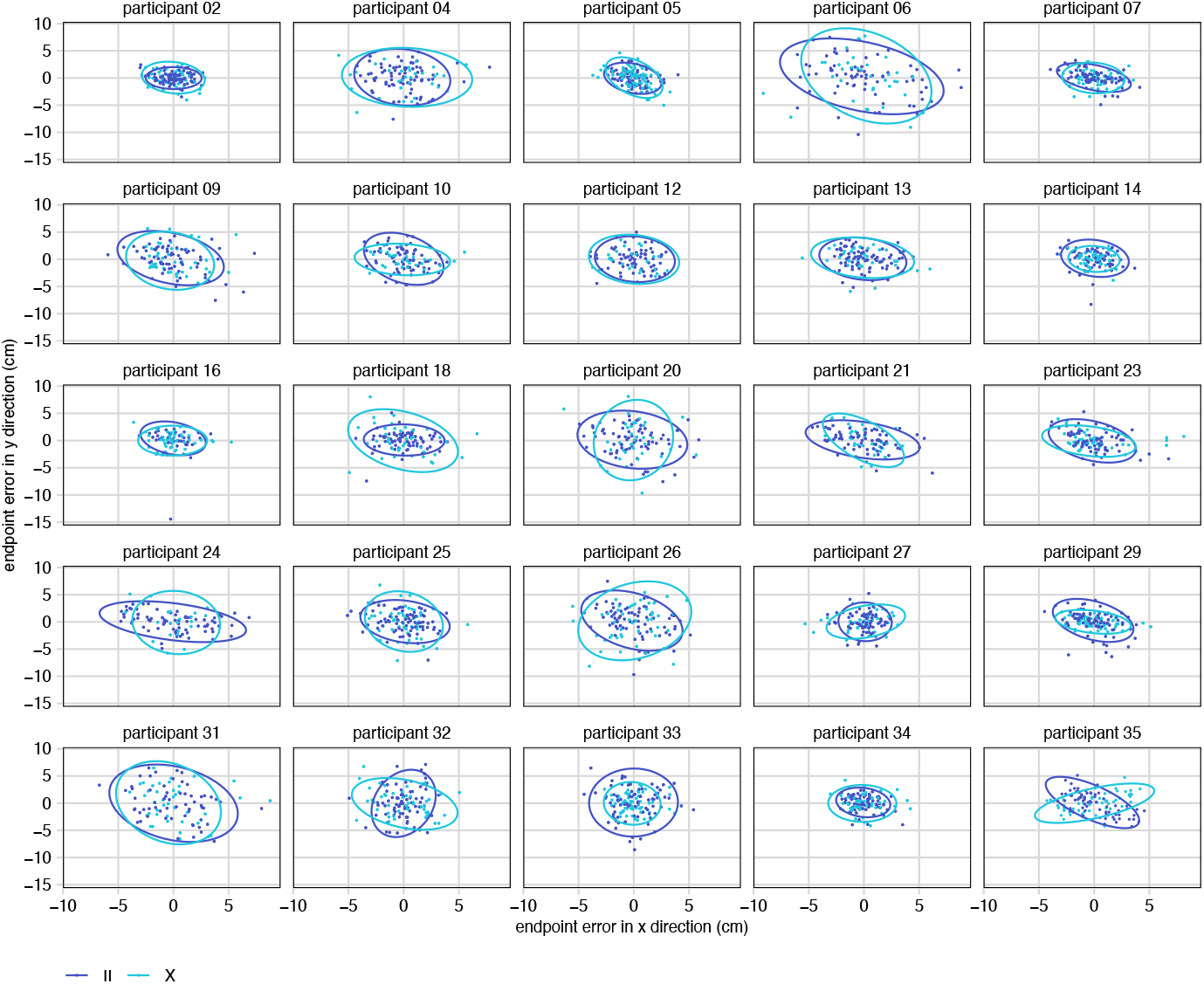
Endpoint variability of mouse click responses to distal stimuli in Experiment 5. We included only trials in which the correct quadrant had been chosen. Left-side and right-side endpoints were merged and are displayed as if all endpoints had ended in the left upper quadrant (see Methods), centered on (0, 0). Ellipses illustrate variability in the uncrossed (dark blue) and crossed (light blue) postures by indicating 95% confidence boundaries of the respective point clouds. A spatial pointing bias towards the opposite side of space in the crossed posture would be evident in a leftward bulge of the light blue data cloud, relative to the dark blue cloud, in a given participant.

#### (Fourth) replication of dependence of performance on number of stimulated limbs

Performance with crossed limbs was again better for stimulus pairs presented to a single limb than for those presented to two limbs, illustrated in Fig. 10e (post-hoc tests: anat:X, z = 7.21, p < 0.001; spatial:X, z = 2.26, p = 0.047). Thus, the advantage of same-limb stimulus pairs occurs independent of the response mode, with button presses, verbal responses, and explicit localization.

## Discussion

We investigated limb assignment in touch using non-spatial comparison judgments (“same limb” vs. “different limbs”), motivated by evidence that humans sometimes confuse on which limb a touch occurred even between distant effectors such as a hand and a foot. Three key findings emerge across all five experiments.

First, Limb crossing robustly impaired limb assignment decisions in all experiments, even when the relevance of spatial stimulus characteristics and spatial response demands were minimized. This impairment generalized across limb sets, affecting arm–leg pairs (Exp. 4) similarly as arm–arm and leg–leg pairs, extending prior reports of categorical hand–foot confusions under crossed postures (Badde et al., 2019; Martel et al., 2022; Schicke & Röder, 2006).

Second, this crossing effect was amplified by external-spatial instructions and reduced by anatomical similarity. Specifically, crossing effects were larger when participants classified pairs by side-of-space rather than by limb identity (Exp. 2, 5), whereas pairs sharing a dermatome yielded smaller crossing effects (Exp. 3). Thus, external framing increased conflict with categorical limb codes, while anatomical features facilitated assignment, consistent with anatomical determinants rather than automatic reliance on external stimulus coordinates. These patterns are incompatible with the long-standing idea that errors in tactile choice tasks derive from a 3D-spatial stimulus code (“space-first”). They converge with the error patterns observed in multi-limb TOJ data showing that limb type, body side, and canonical side—not the touched limb’s current external side—govern misassignments (Badde et al., 2019).

Third, limb assignment errors also manifested in pointing, though the error pattern implied categorical choice errors rather than degrading of continuous localization (Exp. 5). Limb crossing did not significantly modulate endpoint error in pointing responses, suggesting that crossing affected the categorical target selection (which of the four locations relevant in the experiment) more than uncertainty about the stimulus’s 3D location. This dissociation aligns with evidence that stimulus location of TOJ stimuli is constructed post hoc once the limb has been chosen, suggesting that limb choice precedes and determines localization (“limb-first”, Maij et al., 2020). Moreover, errors occurred primarily on distal, crossed portions of the limbs, consistent with the crossed-limb deficit’s spatial specificity to regions distal to the crossing point (Azañón et al., 2016).

Below, we place these three key findings – the relevance of posture for tactile limb assignment, the disadvantage of spatial coding, and the categorical nature of pointing errors – into a larger context, sketch out an integrated theory, and identify research gaps that need filling to further flesh out a firm theoretical explanation for limb assignment errors, crossing effects, and their relation to precise tactile localization for goal-directed actions.

### Touch localization is indirect

It seems a widely held assumption that assigning touch to a body part is based on a straight-forward read-out of somatotopic information contained in S1 maps. However, confusion of hands and feet, as evident in our study and other reports (Badde et al., 2019; Schicke & Röder, 2006), is inconsistent with this assumption (see Kitazawa, 2002, for related arguments).

Multiple experimental findings suggest that tactile-spatial processing relies on identification of the limbs on which the touch occurred (“limb-first”). Perceived tactile distance is systematically distorted by body-part segmentation: identical physical distances are judged longer when they straddle the wrist than when they lie within the hand or within the arm (de Vignemont et al., 2009), showing that continuous skin-space judgments depend on which body part(s) are implicated. Similarly, two tactile stimuli with a fixed distance are perceived as more similar if they both occur on the arm than on arm and hand, as indexed by the mismatch negativity, an event-related brain potential that is sensitive to perceived stimulus similarity (Shen et al., 2018). Finally, the afore-mentioned post-hoc localization of TOJ stimulus location after (not before) a hand has been chosen, too, implies that stimulus localization follows, rather than precedes, limb identification. In sum, the narrative about tactile localization via identity coding in S1 is inconsistent with experimental findings that precise tactile localization involves identifying first which body part received the touch.

### Limb assignment errors are context-dependent

Tactile limb assignment errors are highly task-dependent. For example, crossing effects are more prominent when two stimuli rather than only a single stimulus must be evaluated (Badde et al., 2016, 2019; Kitazawa et al., 2008; Shore et al., 2002), just like in our Experiment 5. More basic still, no one (to our knowledge) has ever reported that participants perceived hand stimuli as having occurred on the feet when the experiment only involved the hands. In the four-limb TOJ task (Badde et al., 2019), misattributions presumably occurred because the four limbs were relevant either as potential stimulus locations or as potential response limbs. In our Experiment 4, participants sometimes reported that an arm and a leg stimulus had occurred on a common limb, just like they reported that stimuli on two homologous limbs belonged to a common limb in our remaining experiments. Thus, the current task appears to create a strict frame for what kinds of conscious percepts can arise, and seemingly absurd confusion of arms and legs occurs only when the task context allows for them.

This context dependence is likely why limb confusions are rarely reported in everyday life. It is hard to say whether limb confusion ever occurs in real-life contexts. For instance, during martial arts sparring, all four limbs are task-relevant, and one might expect that erroneous assignment of tactile input does indeed occur. Yet, given the high confidence about erroneous limb assignment (Badde et al., 2019), they may simply go unnoticed, and negative consequences, such as an inadequate response to an opponent attack, be attributed to inattention or lack of (motor, not perceptual) skill.

### Stimulus choice in the limb assignment task is categorical

However, task context is defined not only by the limbs involved, but likely also by the stimulus set. The present limb assignment task is a choice task, as are many paradigms used in tactile-spatial coding research, such as TOJ tasks. In fact, limb crossing has typically been studied with highly reduced stimulus sets, often only one stimulus per limb. For example, a typical TOJ task involves just two stimuli, one on each hand. Such reduced stimulus sets likely evoke categorical processing strategies. This contrasts with how tactile choice paradigms with crossed limbs have usually been framed, namely assuming that tactile processing necessarily involves precise localization in external space (Azanon et al., 2016; Badde & Heed, 2016; Kitazawa, 2002). Similarly, tactile localization and postural coding are often conceptualized as Bayesian integration of prior experience and current sensory input with a focus on estimating a precise tactile location (Elmas et al., 2025; Goldreich & Tong, 2013; Martel et al., 2022; Peviani, Joosten, et al., 2024; Peviani, Miller, et al., 2024). This focus is misleading in two- or few-alternative forced-choice tasks.

In our study, the stimulus set was also small and involved four locations. Even in Experiment 5, where two stimulators were placed near each ankle and knee to avoid stereotypical pointing, participants may still have treated them as cues for the same body location. We therefore suggest that participants interpreted tactile stimuli as cues for pointing to one of four relevant locations. Similarly, studies requiring reaching (Brandes & Heed, 2015) or eye movements (Overvliet et al., 2011) to tactile stimuli on crossed limbs also involved only two target locations, likely encouraging categorical choices. If crossed postures increased uncertainty about external stimulus location, movements should have shown greater endpoint variability or biases toward the limb’s canonical side. Although these studies focused on trajectories rather than endpoints, their figures do not suggest such endpoint biases – in line with our finding that pointing variability was unaffected by limb crossing in our Experiment 5 (see Fig. 3 in Overvliet et al, 2011; Fig. 2 in Brandes et al., 2015). Instead, trajectories reflected uncertainty about which target to move toward, including delayed trajectory corrections and occasional “turn-arounds.” This pattern suggests that uncertainty affects target selection rather than subsequent spatial computation of the movement endpoint. Note, we do not argue that goal-directed movements proceed without precise localization. Rather, we argue that it is important to differentiate categorical limb choice from precise localization of tactile stimuli, that it is the former that is affected by limb crossing, and that limb choice precedes precise localization and movement planning.

There are, in fact, other pointers in the literature that crossing effects are based on categorical rather than precise spatial information. These findings suggest that participants represent an overall “scene”, or layout, imposed by the task that emphasizes specific spatial aspects but does not involve precise spatial coding. First, in the original report of the TOJ crossing effect (Yamamoto & Kitazawa, 2001), the effect was strongly reduced when reversing the hands did not involve physically crossing the arms but curving one arm around the other hand (see their Fig. 5d). Thus, arm crossing, but not precise hand location, triggered substantial confusion of the two hands. Second, crossing effects remain prominent when both hands are placed in one spatial hemifield (Aglioti et al., 1999; Heed et al., 2016), even though this posture places one hand is in its regular hemifield. Thus, crossing effects appear to depend on the involved limbs’ relative positions to each other, rather than their precise location in space. And third, TOJ for stimuli of the middle and ring finger of one hand are affected by finger crossing just as much as TOJ for stimuli on the two hands by arm crossing, despite practically no spatial distance remaining between the fingertips and the hand being positioned in its regular hemifield–that is, both fingers positioned in their regular hemifield but reversed relative to each other. In sum, limb crossing appears to evoke a categorical, not a spatially graded, change in how tactile stimuli are evaluated in the context of tactile choice tasks.

Our proposal of crossing effects reflecting categorical choice errors within a small set of stimuli inherent in the task makes the testable prediction that crossing errors should be reduced or eliminated if an experiment used a task set that discourages categorical representation of stimuli – a notable gap in the current research landscape.

### Integrating limb choice with the multilateration theory of tactile localization

It has been proposed that tactile localization on the skin exploits multilateration, the localization strategy used in GPS systems (Miller et al., 2022). In multilateration, an object’s location is determined based on its distance to each of several known landmarks–for instance, the location of your car relative to several GPS satellites. According to the model, tactile localization involves determining the location of a touch by multilateration based on its distance to several body landmarks. As an example, the wrist and elbow are plausible landmarks for locations along the lower arm. The 3D location of a touch anywhere on the lower arm could be derived from the 3D position of the landmarks, because the arm is fixed between them. However, localization would not have to be performed in 3D space by default; multilateration could be performed abstractly, for instance in what is sometimes referred to as a “body representation”, independent of how the body is currently positioned.

Notice, that in this scheme the landmarks (i.e., body part) must be chosen first to then allow localization of the tactile stimulus. This requirement introduces the apparent circularity that adequate landmarks must be determined although is not known where the stimulus was: stimulus location appears to be necessary to know which landmarks apply, but landmarks are required to know where the stimulus was in the first place. This conundrum can be resolved by the touch’s association with its canonical space. One can conceptualize the canonical space as a prior not for a precise spatial location but rather for an extended region. Touch is typically perceived together with postural and visual information. Throughout life, an individual experiences touch together with a range of postures, but with much higher probability on the anatomically compatible than the opposite side of space. Accordingly, tactile stimuli on, say, the lower arm become associated with an area of space that may not be precisely defined in size but is clearly lateralized in space – in fact, one might treat this prior as categorical rather than Cartesian (see Fig. 12a). It is this lateralization of the prior we propose guides the choice of the respective landmarks.

**Figure 12:**
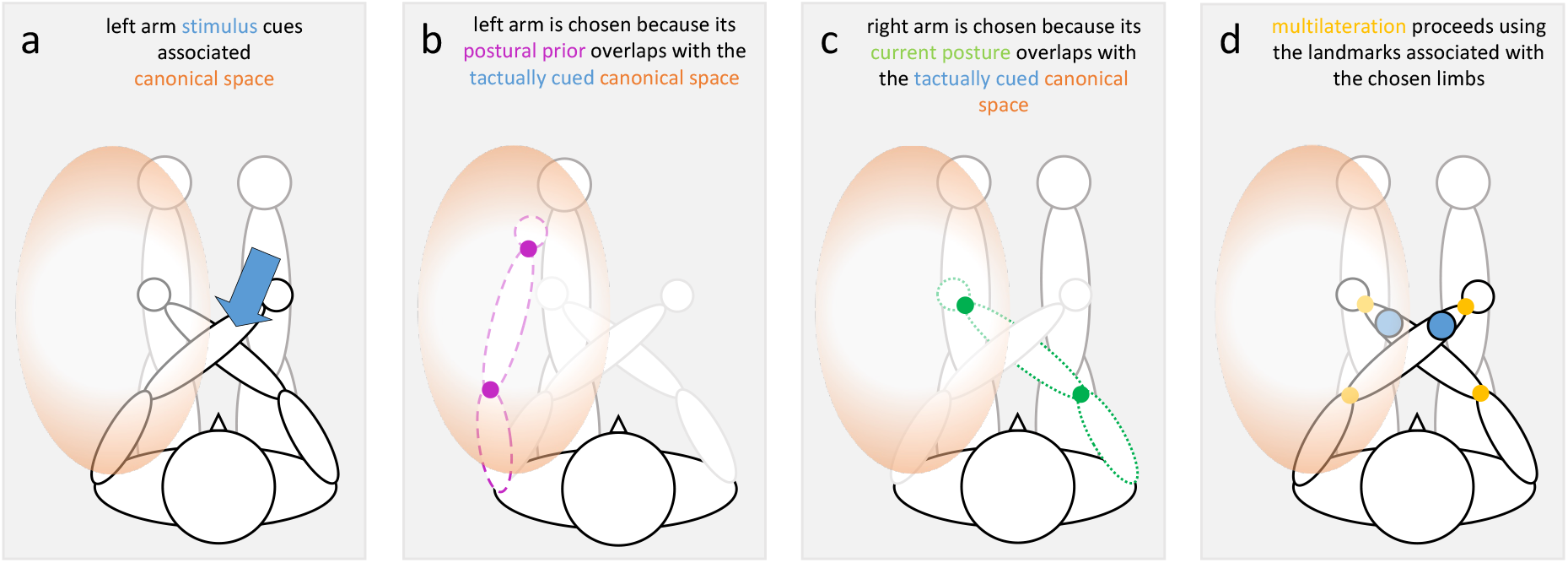
Illustration of the proposed mechanisms involved in limb confusion. In this illustration, a stimulus is presented to the distal location on the left, crossed arm. **(a)** The left arm stimulus (blue arrow) cues the associated canonical space (orange oval); this association has been built up by previous experiences of a touch to this location and the concurrent visual and proprioceptive spatial information. Because actions typically occur on the same side of space as the limb’s bodyside, this prior is strongly lateralized. The oval shape is not meant to depict a presumed true shape of this prior. **(b)** The left arm’s landmarks are chosen because the arm’s postural prior (filled circles on the pink dashed arm) overlaps with the tactually cued canonical space. Like the canonical space, this prior is based on long-term experience, but it relates to body posture, not to the space in which a touch is expected to be located. The faint depiction of the crossed arms illustrates the current posture **(c)** The right arm’s landmarks are chosen because the arm’s current posture (filled circles on the green dotted arm) overlaps with the tactually cued canonical space. **(d)** Multilateration proceeds for each set of landmarks that has been identified as overlapping with the tactually cued canonical space. For the left arm, this results in a cue to the right, distal stimulus location; for the right arm, it results in a cue to the left, distal stimulus location. This latter cue matches the tactually cued canonical space.

This canonical-spatial prior of tactile location is conceptually distinct from the concept of a default posture, which refers to a presumed representation of the limbs’ and body’s “standard” posture (Azañón & Longo, 2019; Tamè et al., 2019). There is in fact evidence for such a postural prior, and both a limb’s current and its default posture are coded spatially (Peviani, Joosten, et al., 2024; Peviani, Miller, et al., 2024). Moreover, participants make distinct localization errors when they localize touch delivered to the fingertips than when the fingertips are cued verbally, supporting the existence of separate priors for tactile location and posture (Elmas et al., 2025).

Limb assignment errors can be explained by connecting multilateration, the canonical-spatial prior, the spatial coding of the limbs’ current and default posture, and the task-dependence of limb confusion. We draw on Badde et al.’s (2019) proposal that phantom errors in their four-limb TOJ study emerged because touch was assigned to an incorrect limb that was currently positioned in the canonical space of the limb that received the first touch. Due to limb crossing, the current posture of the opposite limb overlaps with the canonical space that was cued through the tactile stimulus. For instance, with the arms crossed, a touch to the right arm is associated with the right side of space; this space is, however, currently occupied by the left arm. We propose that this overlap of the touch-associated spatial prior and another limb’s current posture evokes multilateration of the touch on the incorrect limb (see Fig. 12). Moreover, limbs are only considered as having received a touch if they are relevant in the current context (i.e., experiment).

Yet, participants do not err in all trials; in fact, their judgments are often correct. We propose that correct choices involve choosing the landmarks of the correct limb by its prior association with the tactually cued canonical space. In effect, this can be viewed as the interaction of two priors, namely the tactile canonical spatial prior and the limbs’ default posture (see Fig. 12b). Thus, choosing the correct landmarks involves the touch having cued the canonical space, and that space in turn being associated with the limb whose postural position is typically in that space. In other words, according to our proposal, choosing the correct limb’s landmarks has nothing to do with matching the stimulus’s true 3D location with the limb’s current posture. Moreover, and critically, we propose that multiple sets of landmarks can be selected; accordingly, multilateration can be performed both for the limb associated with the tactually cued canonical space and with a limb that currently occupies that space.

But why would the cognitive system allow for multilateration based on different landmark selection criteria – here, landmarks associated either with the canonical space of a touch vs. limbs currently placed in a specific area of space? It has been shown that multilateration can account not only for localization on the arm but also for localization on tools (Miller et al., 2023). Perceiving tactile location on the tool is based on the vibratory patterns evoked by the touch in the tool, sensed by distributed receptors on the hand, whereas perceiving tactile location on the body depends on the identity of the sensors that have registered the touch (Miller et al., 2018). Yet, the processing of tactile location appears to be unified across these different modes of localization very early along the tactile processing hierarchy (Fabio et al., 2024; Miller et al., 2019). Thus, the human cognitive system has evolved to be able to localize touch on non-body objects using many of the processes also used for “regular” tactile processing. For a touch localized on a tool, the touch is registered on the hand but is assigned to a categorically different object – the tool. Thus, the ability of tool use has required dissociating the location of where a touch is registered (hand) from where it is consciously perceived (tool). Similarly, the choice of landmarks (e.g., tip and handle of the tool) for localizing along the tool must dissociate from the anatomical location (hand) at which the tactile input arrives and integrate visual information to interpret the tactile input spatially (Holmes et al., 2007; Iriki et al., 1996; Maravita et al., 2002). It may be for this reason that landmarks for localization can be chosen based on multiple criteria, and touch perceived on a limb other than that which was stimulated, though proof for this conjecture is lacking.

### A proposal for tactile choices as based on multiple categorical features or cues

The last puzzle is that the crossing effect is much more pronounced for distal than proximal stimuli – just as has been reported for TOJ (Azañón et al., 2016). Only distal stimuli changed sides in the crossed posture, because participants crossed their arms over the middle of their forearms or their legs over the middle of their shins. Once again, this finding seems to imply circularity: how would it be possible that confusion occurs mainly for stimuli that changed their side without having localized them first? We propose that this is possible because participants base their judgments on categorical stimulus locations, involving the following steps.

The first step involves only tactile and postural spatial priors. Multilateration of stimulus location is performed on the landmarks associated a priori with the tactually cued canonical space. For example, tactile stimuli on the left crossed arm cue the left side of space (tactile prior); this, in turn, cues using the left arm’s landmarks (postural prior; see Fig. 12b). However, the reduced stimulus set of only 4 possible locations constrains this process to identifying one of the four relevant locations–that is, multilateration is performed only with the precision required in the context of the categorical choice – for instance, by determining whether the stimulus was “distal” or “proximal”.

In a second step, current limb posture is considered (see Fig. 12c). Because the limbs are crossed, part of the incorrect arm intrudes the tactually cued canonical space. This triggers mulitlateration based on the incorrect arm’s landmarks. For distal stimuli, there is now conflict regarding the choice of one of the four possible locations: prior-based processing suggested that touch occurred on the correct arm; however, considering that arm’s current posture now places the identified location (distal on the left arm) outside the tactually cued canonical space. Concurrently, multilateration on the incorrect arm suggests a distal location that matches the cued canonical space. Resolution of this conflict results in increased RT and errors. Presumably, responses are more often correct than not because more cues point to the correct than the incorrect limb; however, conflict resolution may also depend on down-weighting irrelevant (i.e. current posture) information (Badde & Heed, 2016). A weighting account fits well with our finding that anatomical aspects (dermatome, Exp. 3) supported task performance.

For proximal stimuli, no such conflict occurs: after considering current posture, the location identified by multilateration is consistent with that determined via the priors for the touched arm. By contrast, the location determined by multilateration on the incorrect arm is outside the cued canonical space. Accordingly, identification of the correct arm’s proximal location is supported consistently by all sub-processes. Notably, participants made slightly more errors for proximal stimuli on crossed than on uncrossed arms. This supports our proposal that the arm overlapping the stimulus as localized in canonical space indeed evokes projection of the stimulus location on the other arm: in rare cases, the stimulus is then erroneously assigned to the incorrect arm in the crossed posture. Such errors do not occur in the uncrossed posture, because the landmarks of the incorrect arm do not overlap with the cued canonical space, so that multilateration is not triggered for that arm.

Our proposal that multilateration is performed with the landmarks that currently lie in the tactually cued space can also explain the confusion between arm and leg (Experiment 4). When an arm and a leg are involved, the leg’s landmarks can be inserted erroneously, just as the incorrect arm’s landmarks are inserted when only the arms are involved. This confusability may depend on the arms and legs being structurally similar, which allows representing their respective location categories (distal vs. proximal) equivalently. It is of note that some neurological patients either feel a touch applied to one arm also on homologous location of the other arm (termed synchiria), or cannot tell on which of the two limbs the touch occurred (termed allochiria; cf. Medina & Coslett, 2016). It remains to be seen whether these neurological symptoms can be related to the proposed multilateration and limb assignment processes. For instance, it is not clear whether participants perceive correctly and incorrectly assigned stimuli on homologous locations of the limbs. This is because our experiments involve categorical choices rather than precise localization.

Our proposed account also explains why the external-spatial task (Exp. 2 and 5) impeded performance: this is because tactile stimuli are not automatically remapped into 3D space, and so the external location-matching task requires additional, non-default spatial processing to obtain the information required for the spatial version of the task.

### A non-spatial account of tactile coding is consistent with previous findings

We have emphasized here that our experimental paradigm has avoided all possible aspects of space regarding both stimuli and responses. Some previous studies have attempted similar aims with different approaches. One study tested whether stimulus location elicits spatial compatibility effects when it is irrelevant for the task (i.e., a Simon effect). Participants had to judge the intensity of tactile stimuli on the hands as high or low by depressing one of two foot pedals (Medina et al., 2014). When the response involved the foot on the same body side as the stimulated hand, responses were faster – regardless of arm and leg posture. Thus, external stimulus location did not affect responses in this experiment. Another study asked participants to respond to a non-spatial aspect of visual stimuli, namely, color; the visual stimulus was preceded by a task-irrelevant, tactile cue on the hand, with the hand placed either near to, or far from, the visual stimulus location (Azañón et al., 2010). Spatial congruence of the tactile cue with visual stimulus location sped up participants’ responses; thus, the spatial side of the tactile stimulus was considered even though spatial aspects of both touch and visual stimuli were irrelevant to the task. However, visual stimuli inevitably imply a spatial location and, thus, may automatically induce processing of the tactile location when both types of stimuli occur together. This reasoning is supported by the finding that tactile cueing in fact affects visual judgments at the limb’s default location during the first ~150 ms following the tactile cue, and at the external-spatial location only later (Azañón & Soto-Faraco, 2008). This remapping of cue location may well be based on hand location rather than stimulus location (cf. Badde & Heed, 2023; Klautke et al., 2023), given that transcranial magnetic stimulation (TMS) of parietal cortex disrupts matching a tactile stimulus on a hand with a visual location (Bolognini & Maravita, 2007) as well as matching the spatial elevation of tactile locations on the arm and face (Azanon et al., 2010) while leaving proprioception and tactile detection unaffected.

Overall, our proposal is consistent with previous findings, even if these have typically been interpreted to support external-spatial coding of touch. In turn, it is plausible that also other crossing effects, such as that observed in the TOJ task, reflect errors in assigning a tactile stimulus to a body part and not errors in determining stimulus location in space (Badde et al., 2019; Maij et al., 2020).

### Summary and outlook

In summary, we have presented evidence suggesting that the prevalent idea that tactile location is automatically remapped into a 3D-spatial code is likely false and have laid out an alternative theoretical account. Previous work has often leaned heavily on tactile crossing effects as a presumable marker for automatic 3D-spatial recoding of tactile location. We argue here that crossing effects arise from overlap of the touched limb’s default spatial position and the position of other, currently relevant limbs, leading to conflict between the different available choice options in tasks both with spatially specified responses such as button presses with the stimulated limb, and with non-spatial responses, such as verbal report and responses with limbs that do not receive stimulation.

Our experiments here used tactile choice paradigms in which only few spatial locations are stimulated during the entire experiment and responses are categorical. It is an open question whether the absence of automatic 3D spatial coding of tactile stimulus location is unique to such choice tasks, or whether it extends to paradigms in which a precise, localized response is required, such as when participants must touch a tactually stimulated location with their index finger, or when stimuli (appear to) move across the skin, such as in the rabbit illusion, with which began our introduction. In sensorimotor neuroscience, it is an established idea that (visual) stimulus and effector locations are coded in a vision-based, eye-centered code (Andersen & Cui, 2009). Based on this framework, as well as based on findings that imply spatial matching of visual and tactile spatial location in multisensory contexts, research on tactile-spatial processing has been based on the idea that touch is automatically recoded in a 3D spatial, likely visual, code. When a movement must be planned, it would then be irrelevant whether the target was presented visually or tactually, because the common 3D spatial code that is available across modalities would support movements independent of the sensory input modality. In contrast to this reasoning, we have argued that the 3D location of touch is inferred indirectly, for instance, by multilateration (Miller et al., 2022), and related to space via the current, posture-dependent location of the involved body landmarks. In this framework, movement towards touch is localized locally on the limb relative to landmarks such as the joints, and the movement planned towards the spatial location of the touched limb (cf. Badde & Heed, 2023; Klautke et al., 2023). Thus, the coding principles we have uncovered here for choice tasks may be universal across all tactile contexts – but only future research will show whether this proposal holds.

## Methods

### Data and code availability and preregistration

All single trial data are freely available at the Austrian NeuroCloud (ANC, https://anc.plus.ac.at):

- Experiment 1: Heed & Fuchs, 2025, https://doi.org/10.60817/9fzf-v802;
- Experiment 2: Heed, Fuchs, Burbach, & Rödenbeck, 2025, https://doi.org/10.60817/qjdm-g591;
- Experiment 3: Heed & Fuchs, 2025, https://doi.org/10.60817/b64r-9370;
- Experiment 4: Pollner, Heed, & Fuchs, https://doi.org/10.60817/xwj2-j822;
- Experiment 5: Heed, Fuchs, & Weiner, https://doi.org/10.60817/f8yj-bg69).

These data include participants that we removed (if any), or considered removing, from our analysis, thus allowing re-running our analyses with and without these participants’ data. The commented R code used for analysis and figure generation of all five experiments is available at the Open Science Framework (OSF, https://doi.org/10.17605/OSF.IO/ZH5RG). Experiments 4 and 5 were preregistered at OSF (Experiment 4: https://osf.io/z4d3r; Experiment 5: https://osf.io/xdhu6).

### Experiment 1

#### Determination of sample size and sensitivity analysis

Crossing effects are very prominent in many paradigms (Badde & Heed, 2016; Heed & Azañón, 2014) and have been found to be highly significant with very small sample sizes such as 10 (Heed & Azañón, 2014; Röder et al., 2004; Schicke & Röder, 2006). Because the current paradigm was new, we aimed at recruiting roughly twice as many participants in this and the following experiments.

#### Participants

Twenty-one participants took part in Experiment 1. They were selected to be between 18 and 40 years old, right-handed, have normal or corrected-to-normal vision, and be free of neurological disorders as well as tactile and auditory deficits by self-report. Nine participants reported themselves as female, 9 as male. They were 19-31 years old (mean: 23.0). Participants gave their written consent to participate in the study after they had been fully informed about all details of the experiment. They received course credit for their participation. All studies we report were approved by Bielefeld University’s ethics committee (file number 2017-114).

#### Setup

Three electromagnetic solenoid-type tactile stimulators with 1.8 cm diameter (Dancer Design, St. Helens, UK) were attached to each arm with adhesive rings. For stimulator placement, the participant’s lower arm was measured from the wrist to the elbow crease, and the upper arm from the elbow crease to the highest point of the shoulder. The first stimulator was attached near the wrist, at 1/6 of the participant’s lower arm length. The second stimulator was attached on the lower arm at 1/6 of lower arm length from the elbow. The third stimulator was attached on the upper arm at 1/3 of the upper arm’s length from the elbow.

Tactile stimuli were 15 ms long 200 Hz vibrations. They were controlled and amplified for stimulator output by a custom-built control unit (Neurocore, Hamburg, Germany) that was connected to a Windows 7 personal computer via the universal serial bus (USB). Stimulus intensity varied slightly from trial to trial, with each stimulus being presented with one of three intensities (a standard current output directed to the stimulator, +/-5%) chosen randomly, with equal probability, from trial to trial, and independently for each stimulus. This manipulation prevented participants from using idiosyncratic characteristics of the stimuli to make their judgments, for instance that one stimulus of an evaluated pair was always perceived as weaker than the other even though stimulator output was identical. All stimulus intensities were clearly supra-threshold. Participants wore ear protection to shield any sounds the tactile stimulators made. They had their eyes open and fixated a cross about 80 cm straight ahead at eye level.

Participants gave responses by lifting the toes or heels of both feet, respectively. The feet rested on response buttons which registered a response when the respective part of the foot was lifted. We used as reaction time the time interval between the start time of the second stimulus and the registration of the first of the two feet’s button responses.

The experiment was programmed with PsychoPy (v1.85.1, Peirce et al., 2019).

#### Trial structure

Each trial started with a short tone, presented via headphones, marking the beginning of a trial. To eliminate any possibility of anticipation effects of the tone on subsequent events, the tone was followed by a random interval lasting between 500 and 1000 ms. Next, two tactile stimuli were presented. The participant had to respond whether the two stimuli had occurred on one or on two limbs. These two responses were mapped to lifting toes or heels, and this assignment was counterbalanced across participants. Each response was acknowledged by a short beep, independent of whether it was correct or not, followed by a fixed intertrial interval of 1000 ms. If participants responded before the second stimulus had been presented, or if no response was registered within 3000 ms, the trial was marked as invalid and repeated at the end of the respective block. If only a single foot’s (rather than both feet’s) response was registered, the trial was also considered invalid and repeated at the end of the block. Invalid trials were indicated to the participant by a higher beep than the regular response acknowledgement.

#### Experimental Design

The experiment comprised four factors. First, the arms were placed either uncrossed or crossed (factor: Posture; levels: uncrossed vs. crossed). Participants first performed the experiment in one and then in the other posture. We counterbalanced across participants which posture came first and which arm was placed above the other. When the arms were uncrossed, they were held straight, so that the three stimulators were arranged on a straight line away from the body. When the arms were crossed, one lower arm was placed over the other at about the lower arm’s middle. Participants placed their arms so that the two upper stimulators of the left arm, together with the lower stimulator of the right arm, formed a straight line away from the body, and vice versa. Thus, the stimulus layout was roughly identical in uncrossed and crossed postures.

Second, the two stimuli were either presented on the same part of the arm, or on different parts of the arm (factor: Number of Segments; levels: one vs. two). In each trial, one stimulus was always on the low position near the wrist. The other stimulus could be presented also on the lower arm, so that the two stimuli were both presented on the same body segment. Alternatively, the other stimulus was on the upper arm, so that the two stimuli were distributed across lower and upper arms. The order of the two stimuli was counterbalanced across the experiment, with the wrist position occurring first or second. All analyses collapsed across these two trial types.

Third, the two stimuli were presented either on the same limb or on opposite limbs (factor: Number of Limbs, levels: one vs. two). Thus, stimuli could both be presented on the same arm segment (lower arm), but on two arms.

Fourth, the two stimuli followed each other at four different stimulus onset asynchronies (SOAs): 50, 100, 300, or 800 ms. We included this factor to test whether task performance would level at large SOAs. In TOJ tasks, participants are usually significantly better in the crossed hands posture when the SOA is > ~300 ms than when it is shorter and aimed to test whether a similar effect would be evident when limb identity must be evaluated.

#### Procedure

We ascertained that all stimuli were clearly suprathreshold. Participants performed a short practice of 16 trials in uncrossed and crossed postures. Each factor combination was repeated 48 times, resulting in 1536 trials in total. Trials were distributed across 16 blocks of 96 trials each. Participants could rest after each block and took a break of at least 30 minutes after half of the experiment. The experiment took about 3 hours including instructions, setup, and breaks.

#### Analysis

We analyzed data with R (version 4.6.1, R Core Team, https://www.R-project.org) in the RStudio programming environment (PBC, Boston, MA, USA, http://www.rstudio.com/), using packages tidyverse (Wickham et al., 2019), magrittr (Bache & Wickham, 2022), and afex (Singmann et al., 2023) and prepared figures with ggplot2 (Wickham, 2009) and ggrain (Allen et al., 2021). We report the Balanced Integration Score (BIS), a measure that integrates normalized reaction time and accuracy that is reportedly superior to other integrative RT/accuracy measures (Liesefeld & Janczyk, 2019). Given that we are interested in within-participant differences, we normalized within each participant rather than across the entire sample (Liesefeld & Janczyk, 2019, 2022). Our qualitative conclusions are supported by analysis of RT, accuracy, and BIS alike. RT and accuracy are presented separately in the supplementary information. We used ANOVA as our statistical model. We report both a full model that includes SOA in the supplementary information; however, all effects were consistent across the tested SOAs, and so we dropped the factor from the ANOVA reported in the main text. We performed condition-specific post-hoc tests with paired t-tests, correcting for multiple comparisons using the false-discovery rate (FDR), as implemented in the R package emmeans (Lenth et al., 2023).

#### Selection of participants for analysis

We considered removing two participants for whom our lab notes stated edginess or tiredness; results regarding statistical significance did not change qualitatively whether these participants were included or not and we therefore kept them in the analysis. For a third participant, the data are 39 of the 1536 intended trials short, resulting from a technical error at the end of one experimental block. The trial numbers per cell for that participant are 88-95 of the intended 96. We consider these numbers sufficient to include the participant in the analysis.

### Experiment 2

Many aspects of Experiment 2 were identical to those of Experiments 1. We report here only those experimental details that were different.

#### Participants

We recruited 27 new participants according to the same criteria as in the previous experiments but did not exclude left-handers. The higher number of participants was due to our aiming at 20 (as in Experiment 1) but running the experiment in the context of student teaching, which afforded additional acquisition slots. Fourteen participants reported to be female, 12 male, and one diverse; they were 19-37 years old (mean: 24.7); 26 participants were right-handed and one left-handed.

#### Setup

The general setup resembled that of Experiment 1 but we used a different amplifier for the tactile stimulators. Instead of vibratory stimuli, we used “tap”-stimuli, produced using 25-ms rectangle waveforms. The stimuli were generated using a USB-connected low-latency device (Labjack T4, LabJack Corporation, Lakewood, USA) sending the waveform via the analogue out port to a tactile amplifier (Tactamp, Dancer Design, St. Helens, UK) and then routing the amplified signal to one of the 4 stimulators (same product as in Experiment 1) using a switchbox (Dancer Design, St. Helens, UK). The Labjack’s digital ports were used to control the switchbox and record the responses from the response buttons. A self-programmed library for the Python programming language, “pytact” (https://github.com/xaverfuchs/pytact), provided the functions to present the stimuli and record the button responses.

#### Trial Structure

All details were as in Experiment 1, except that the maximum response time for the participants was decreased to 2500 ms to reduce the duration of the experiment.

#### Experimental design

The experiment comprised three factors. First, each participant performed the experiment with two different response requirements (factor: Instruction, levels: anatomical vs. spatial). Under anatomical instructions, participants responded in the same way as in Experiment 1, that is, whether the two stimuli of a given trial had occurred on the same limb or not. Under spatial instructions, participants instead responded whether the two stimuli had occurred on the same side of space or not. The two instructions were run one after the other, with all other factors nested within Instruction. Instruction order was balanced across participants.

Second, the arms were positioned either uncrossed or crossed (factor: Posture; levels: uncrossed vs. crossed), which was varied between consecutive experimental blocks, with the posture at the experiment’s beginning balanced across participants.

Third, the two stimuli were presented either on the same arm or on opposite arms (factor: Number of Limbs, levels: one vs. two).

In Experiment 1, the shortest SOA of 50 ms had elicited the largest differences between experimental conditions. Other SOAs had elicited similar but smaller effects. Therefore, we dropped the factor SOA in Experiment 2 and used only an SOA of 50 ms.

#### Procedure

We repeated each factor combination 120 times, resulting in a total of 960 trials. We presented the experiment in blocks of 40. In one block, there were 10 repetitions of the four possible stimulus combinations (e.g., left distal and left proximal stimulus), with the presentation order balanced among them. The experiment took about 2 hours including instructions, setup, and frequent breaks.

#### Analysis

We analyzed Experiment 2 in the same way as Experiment 1 with the factors specified above.

#### Selection of participants for analysis

Five of our 27 participants performed around chance in the crossed conditions on average. We ran analyses with and without these participants; excluding them slightly improved some of the statistical comparisons but did not change the overall qualitative result pattern. We therefore report on the full dataset.

### Experiment 3

Many aspects of Experiment 3 were identical to those of Experiments 1 and 2. We report here only those experimental details that were different. We ran Experiment 3 twice, referred to as 3a and 3b; the two instances differed only in the way in which we determined the stimulus locations. We report statistics for the pooled sample.

#### Participants

We recruited new participants who had not participated in any of the other experiments for both Experiments 3a and 3b according to the same criteria as in the previous experiments but did not exclude left-handers.

##### Experiment 3a

Of 20 recruited participants, we excluded one from analysis (see below). The remaining 19 participants (12 female, 7 male) were aged 20-25 years (mean: 21.7). Three of them were left-handed.

##### Experiment 3b

Of 20 recruited participants, we excluded one from analysis (see below). The remaining 19 participants (8 female, 11 male) were 19-35 years old (mean: 23.1). Four of them were left-handed. Participants did not receive course credit in this experiment.

##### Pooled sample

Together, the sample consisted of 38 participants aged 19-35 (mean: 22.4) years.

#### Setup of Experiment 3a

The technical setup was identical to the one used in Experiment 1. Participants sat on a height-adjustable chair. Its height was adjusted so that participants could place their feet on the floor with crossed legs. When the legs were uncrossed, they were placed parallel at hip-width distance, with the feet placed firmly on the floor. When the legs were crossed, one lower leg was placed in front of the other at the shin. In this posture, the feet rested on their outer sides. The low-position stimulators were placed in the opposite-than-normal hemifield and did not touch each other or the other leg. Which leg was placed before the other was kept constant within each participant and balanced across the sample.

We guided stimulator placement according to published borders of the human dermatomes (Keegan & Garrett, 1948; Lee et al., 2008). Dermatome borders vary between individuals, and there is some overlap between dermatomes (Ladak et al., 2014; Lee et al., 2008). Thus, placement may not have been in the targeted dermatomes for some participants. However, testing dermatomes would involve anaesthesia of spine segments, which is an invasive procedure that we did not consider ethically adequate for our study. We placed three stimulators on each lower leg. One stimulator was placed 3 cm above the palpated upper end of the outer leg’s ankle but shifted towards the back of the leg by about 45°; this position presumably belongs to dermatome S-1 (Keegan & Garrett, 1948; Lee et al., 2008). The second and third stimulators were placed ~5 cm below the hollow of the knee, both at an identical distance from the floor when the participant placed their feet firmly on the floor in a normal, seated posture. We measured the leg’s circumference at this height and placed the second stimulator one third of the circumference towards the leg’s outer side, measured from the anterior edge of the tibia. This placed the second stimulator within dermatome S-1 (Keegan & Garrett, 1948; Lee et al., 2008). The third stimulator was placed 20% of the circumference towards the leg’s inner side, placing this stimulator within dermatome L-3 (Keegan & Garrett, 1948) or L-3/L-4 (Lee et al., 2008). Equivalently to Experiments 1 and 2, a given trial’s stimulus pair always comprised the distal ankle stimulus plus a proximal knee stimulus. Consequently, there were stimulus pairs that shared the S-1 dermatome and pairs that belonged to different dermatomes.

Responses were equivalent to those of Experiment 1 but executed with the two hands rather than the feet. A 20 cm-wide shelf was placed in front of the torso, and four round, 7 cm diameter response buttons were placed on it in a rectangular layout; left and right buttons were 15 cm apart measured from their inner sides, and the buttons in the front and back were separated by 3 cm. Each hand rested on one pair of buttons, with the fingers depressing the respective front button and the hand bale the back button. Participants lifted either all 10 fingers to release the two buttons in the front, or they lifted the two hands’ bales to release the two buttons in the back, resulting in two possible response alternatives. A fixation cross was present ~130 cm from the participant’s head.

#### Setup of Experiment 3b

Here, the upper stimulators were attached to the upper leg. Participants sat on the chair and rested their feet on a small, low table with their feet stretched out and, if required for comfort, the knees slightly bent.

One reason for the uncertain mapping of skin to dermatomes is that each nerve from the leg feeds into several spinal segments within the lumbosacral complex. We chose regions such that one pair of lower/upper leg locations reliably fell into S-2, and the other upper leg stimulus into L-2 (Lee et al., 2008). These dermatomes have no overlap in the spinal segments at which they enter the lumbosacral complex (Schünke et al., 2018a, 2018b), making the anatomical separation of the respective stimulus pairs even more credible than in Experiment 3a. The lower leg location was on the back of the leg, 2 cm above the ankles, half-way medial of the posterior lower leg in the innervation area of the sural nerve, which is a branch of the sciatic nerve (spinal segments: L-4 to S-2, mainly S-2). We placed the dermatome-matched, upper leg stimulator on the back of the thigh, about one third of upper-leg length above the hollow of the knee, in the innervation area of posterior femoral cutaneous nerve (spinal segments: S-1 to S-3, mainly S-2). We placed the non-matching dermatome stimulator on the dorsal thigh at half the length from knee to hip in the innervation area of the lateral cutaneous nerve (spinal segments: L-2 to L-3, mainly L-2). Thus, the entry points for the two upper-leg locations were clearly separated, with one entering the spine in the sacral, and the other in the lumbal section of the spine, separated by S-3, which is not an entry point for either skin location.

The fixation cross was about 125 cm away from the participants’ head. The response buttons were placed on a tray that participants rested on their lap. An additional break was inserted after 55 of 95 trials in each block to allow participants to relax, because pretesting had shown that the legs could become numb in this experiment’s posture, with one leg resting on the other.

#### Trial Structure

All details were identical in Experiments 3a and 3b. They matched those of Experiment 1.

#### Experimental design

The experiment comprised three factors. First, the legs were positioned either uncrossed or crossed (factor: leg posture; levels: uncrossed vs. crossed), acquired one after another, as in Experiments 1 and 2.

Second, the two stimuli were either presented in the same dermatome or in different dermatomes (factor: Number of Dermatomes; levels: one vs. two). The order of the two stimuli – proximal or distal presented first – was counterbalanced across the experiment. All analyses collapsed across stimulus order. We note that Experiment 1 had not tested a potential modulation by dermatome, despite presenting stimuli on different arm segments. This is because the dermatomes on the arm are not organized according to the upper vs. lower arm but, instead, run in stripes along the entire arms and cover both the upper and lower arms, so that the stimuli of the 1 vs. 2 segments manipulation in Experiment 1 lay in the same dermatome.

Third, the two stimuli were presented either on the same leg or on opposite legs (factor: stimulated limbs, levels: one vs. two). Thus, stimuli could both be presented in equivalent dermatomes on two legs, which both enter the spine at the same segment but, of course, nonetheless belong to different skin regions.

As in Experiment 2, the SOA between the two tactile stimuli of a given trial was always 50 ms.

#### Procedure

The experiment had 4 blocks of 96 trials each, resulting in a total of 384 trials.

#### Analysis

We analyzed Experiment 3 in the same way as Experiment 2.

#### Selection of participants for analysis

We removed 2 participants from analysis – one each from Experiments 3a and 3b. Both had responded 100% incorrect for one single stimulus pair. Each stimulus was part of several stimulus pairs; it is unclear why these two participants responded so consistently incorrect to a single stimulus combination but not to other pairs involving one of the involved stimuli. Anecdotally, stimulation on the upper leg felt unusual to some participants and some reported finding the task difficult for that reason. Still, percentage correct was >50% for most stimulus pairs across participants. Removing the 2 participants in Experiment 3 did not affect the qualitative data pattern and the statistical conclusions drawn from our models.

### Experiment 4

Many aspects of Experiment 4 were identical to those of Experiments 1-3. We report here only those experimental details that were different. Experiment 4 was preregistered at https://osf.io/z4d3r.

#### Participants

Based on a power analysis, we aimed to recruit 30 new participants according to the same criteria as in the previous experiments but did not exclude left-handers. We replaced one participant for whom data had not been properly saved. We excluded 2 participants from the final sample (see below). Of the remaining 28 participants, 9 reported to be female and 19 male, they were 18-31 years old (mean: 23.7); 24 participants were right-handed and 4 left-handed.

#### Power analysis

We used data from experiment for simulation-based power estimation. The code and results of the power estimation are available in the preregistration on OSF. According to the simulations, acquiring data from thirty participants was sufficient to detect differences between the levels of the factor of interest (Limb Set, see below) even if they differed by only 20%.

#### Setup

The general setup resembled that of Experiment 2, but we used a different interface for producing the stimuli. The TactAmp was triggered by a low-latency PCI-port connected data acquisition card (NI PCI-622, National Instruments, Austin, TX, USA). As before the tactile stimuli consisted of a 100 Hz vibrotactile stimulus with three pulses. The two stimuli of one trial always had different intensities: one stimulus was presented at full intensity, the other one at 50% intensity (balanced across all combinations of used locations). The SOA of the two stimuli was 100 ms. Responses were registered verbally: at the start of the experiment, the participants were instructed to respond “tipp” or “topp” depending on whether the stimuli were perceived on the same or on different limbs (coding of the words was counterbalanced across participants). Participants had to respond within 3000 ms. The verbal responses were recorded continuously in the free recording software Audacity (audacityteam.org) using standard vocal microphone connected to a USB audio interface (Scarlett 2^nd^ generation, Focusrite, High Wycombe, UK). To register the sections of the audio stream with vocal responses, brief pulses from the NI card that were send simultaneously with the stimuli to the audio device and recorded on the channel of the stereo recording that was not used for the verbal responses. These pulses were later used to slice the audio recording into segments containing the responses to the stimuli. Responses were decoded from the audio files using a freely available verbal model (vosk; https://alphacephei.com/vosk/) controlled via a custom python script that extracted the responded word verbal reaction time.

#### Trial Structure

All details were as in Experiment 1, except that the maximum response time was decreased to 2500 ms to reduce the duration of the experiment.

#### Experimental design

The experiment comprised three factors. First, each participant performed the experiment with two different limb sets (factor: Limb Set, levels: arm-arm vs. arm-leg). For each participant, the arm-leg condition comprised either the left arm and right leg, or vice versa, distributed evenly across the sample.

Second, the limbs were positioned either uncrossed or crossed (factor: Posture; levels: uncrossed vs. crossed), which was varied between consecutive experimental blocks, with the posture at the experiment’s beginning balanced across participants.

Third, the two stimuli were presented either on the same leg or on opposite legs (factor: Number of Limbs, levels: one vs. two). The limb sets were run in blocks with the block order balanced across participants.

#### Procedure

We repeated each stimulus pair 10 times. We had defined stimulus pairs to present either the distal or the proximal stimulus first. After the data had been acquired, we noticed that one stimulus pair had been coded incorrectly, so that one proximal-first stimulus pair was replaced by a meaningless stimulus combination to all participants. We removed all proximal-first trials from the dataset to obtain a fully balanced trial set. Comparison of full and selected data for the correctly acquired conditions showed that halving the data changed the averaged results for both single participants and the group only minimally. The open data on OSF contain all acquired trials for scrutinization and code to run the paper’s analyses with and without removing data. Because each experimental condition has a mirror-twin (e.g., “same arm” = proximal-right/distal-right plus proximal-left/distal-left trials), each factor combination contained 20 stimuli per participant. Because the voice key responses were analyzed offline, we could not repeat trials in which no response or no clearly identifiable response had been given. We removed such trials from the data, resulting in loss of 109 (2.2%) trials.

The experiment took about 1.5 hours including instructions, setup, and frequent breaks.

#### Analysis

We analyzed Experiment 4 in the same way as Experiment 2, but with the factors specified above.

#### Selection of participants for analysis

We excluded 3 participants from analysis. For one participant, half of the logfile was missing for unknown reasons. This participant was replaced because the error became evident while data collection was ongoing. One participant exhibited chance performance selectively for two stimulus pairs, and one participant for a single stimulus pair. The open data, available at OSF, include all 31 participants (30 recruiting aim, 1 replacement) for scrutinization.

### Experiment 5

Experiment was preregistered at https://osf.io/xdhu6.

#### Participants

We aimed to recruit 30 new participants according to the same criteria as in the previous experiments; the factorial structure was similar to Experiment 4, so we relied on the power analysis of that experiment also for the current one. As is evident from Fig. 11, data files are numbered up to 35. The first participant was excluded because of a programming error that became apparent after acquisition. One participant was excluded due to a technical error in one stimulator which became apparent only some time into acquisition. Three participants did not show (but had received numbers in our acquisition scheme). Thus, we acquired 30 full data sets. We excluded 4 of them (see below); of the remaining 26 participants, 24 reported to be female and 2 male; they were 18-33 years old (mean: 21.4); All participants were right-handed.

#### Setup

Stimulation employed the same technical setup as in Experiment 4. The stimuli were also identical, except that the SOA for the two stimuli was 0 ms. Thus, in the two-stimulus variant of the task, both stimuli were presented simultaneously to avoid localization biases arising from stimulus presentation order. To encourage true localization of the two stimuli and avoid stereotyped responses, two stimulators were attached at each of four locations on the lower legs (near the ankles and knees), resulting in a total of eight stimulators. A monitor (iiyama ProLite T2753MSC, iiyama, Tokyo, Japan; 60 × 34 cm, 1920 × 1080 pixels, 60 Hz), mounted horizontally on an aluminium frame, was positioned over the participant’s hand, with the long side extending away from the body. Participants localized the stimuli by clicking with the mouse cursor on corresponding locations on the monitor. To prevent localization responses from being biased by the cursor position of the previous response, each localization trial began with the cursor appearing at one of two randomly selected start positions. These start positions were centered in the near–far dimension and located 15 cm either to the left or right of the monitor center. The cursor was displayed as a blue circle (diameter: 0.5 cm) for the first localization and as a red circle for the second localization. The first cursor appeared after stimulus presentation and disappeared when participants confirmed the selected location by pressing the right mouse button. The second cursor then appeared, allowing participants to indicate the location of the second stimulus.

#### Trial Structure

The two tactile stimuli were presented concurrently. There was always one proximal, near-the-knee stimulus and one distal, near-the-ankle stimulus. Participants then clicked on the screen to localize them as precisely as possible with a time limit of 3 s. We introduced this time limit as time pressure to elicit erroneous responses. As participants had to indicated 2 locations, we did not expect RT to be a useful measure in this experiment, and so we deemed it important to set conditions in a way to bias responses to be fast and incorrect, rather than slow and correct. Participants chose freely which of the two stimuli to report first. If the time limit of 3 s for a given response had passed, a message (“too slow”) occurred on the screen to encourage participants to respond faster; however, the trial continued until the responses had both been given.

#### Experimental design

The experiment comprised three factors. First, each participant responded in two different ways across different blocks. In the first response mode, there was a photo of a pair of legs, and participants indicated the precise locations at which they had perceived the two stimuli on the legs by clicking on the them with the mouse cursor (“anatomical response mode”). We ran this response mode with the photo compatible or incompatible with respect to the participant’s true leg posture, but we report here only on the compatible condition. As the second response mode, participants indicated the spatial locations of where they had felt the tactile stimuli by clicking onto the empty screen. We instructed them to imagine the screen to be a window through the table surface from straight above, so that they would see their legs through this window and click on the place of the window where they would “see” the stimuli. Thus, the factor Response Mode had the two levels “anatomical” and “spatial”. The response modes were devised in randomized order across participants; the uncrossed and crossed posture was acquired in the same order for a given participant, so that they frequently uncrossed their legs to avoid pain and numbness. Whether a participant first performed the uncrossed or crossed posture was randomized across participants.

The second and third factors were identical to those of the previous experiments: the limbs were positioned either uncrossed or crossed (factor: Posture; levels: uncrossed vs. crossed), and the two stimuli were presented either on the same leg or on opposite legs (factor: Number of Limbs, levels: one vs. two).

#### Procedure

We repeated every factor combination of locations 20 times (e.g., left distal and left proximal stimulus), distributed across 5 blocks following upon each other, resulting in a total of 320 trials. For the spatial localization mode, we also assessed localization of single stimuli. Every location was stimulated 10 times and participants localized the stimuli in the same way as before by indicating the location with the mouse cursor.

The experiment took about 1.5 hours including instructions, setup, frequent breaks, and the non-reported incongruent response mode.

#### Analysis

We used two complementary analysis strategies for Experiment 5. First, we classified each click into one of four quadrants (proximal vs. distal limb location x left or right side). This allowed us to match whether a given click reported by the participant matched one of the trial’s stimuli: for instance, when we had stimulated the right knee, then one of the clicks should have occurred in the lower, right side of the photo or screen. Classification was based on clustering performed over click locations, derived separately by participant, response mode, and posture. We then derived an accuracy measure that was equivalent to the “same/different” response in Experiments 1-4: we classified trials as correct (each click could be matched to one stimulus) or incorrect (at least one click could not be matched). We report only accuracy, because participants had to click twice per trial, prohibiting adequate analysis of RT and, consequently, computation of BIS scores. We employed GLMMs with a logistic link function for statistical analysis because ANOVAs are not strictly adequate for modelling accuracy data (Jaeger, 2008). We used the maximal random effects structure for which models converged (Barr, 2013); this was a full random structure including all interactions, but without correlations between the random predictors (coded as “||” in the model’s random effects specification). As in all other experiments, we followed up on significant interactions with contrasts computed via the emmeans package.

Second, we analyzed the locations reported by participants in the 2D space of the screen and tested whether they exhibit biases towards the opposite side of space. We pooled over the two locations at each body location; e.g., we treated the two stimulus locations on the left ankle as a single location. The two locations were only few cm apart on the skin for all 4 body locations; moreover, one location was always more lateral than the other, so that their distance in the 2D plane of the monitor, or their location on the legs depicted on the screen, was very small. Pilot testing revealed that the endpoint variability for each location was much larger than the variability induced by using two close-by locations. By pooling over the two near-by locations, we thus doubled the number of trials per condition while inducing only minimally more variance than if we had analyzed each location separately. We de-meaned left-side and right-side endpoints separately, then flipped right-side endpoints with respect to left and right, and then merged localization responses of on the left and right sides. Thus, localization data are presented as if all stimuli hat occurred on the left side of the screen.

One participant (participant 30) adapted their crossed posture while performing the spatial response mode. Consequently, they pointed to different locations for some stimuli before and after adaption, which led to inflated endpoint variability for this participant. They were therefore excluded in the endpoint analysis (see Fig. 11, which accordingly shows only 25 participants).

#### Selection of participants for analysis

We excluded 4 participants from analysis. One participant had about 40% localization errors in the uncrossed posture, and three others localized the two stimuli of a trial at the same elevation (i.e., both as proximal or both as distal) in 20% or more trials, although stimuli were always one at a distal and one at a proximal location. The open data, available at OSF, include all 30 participants for scrutinization.

## Acknowledgements

We thank Stefanie Queren, Nadine Krautzer, Georg Pollner, and Paul Weiner for help with data acquisition. This work was supported by the German Research Foundation (DFG) through an Emmy Noether grant to TH (He 6368/1-3).

## Author Contributions

Contributions according to https://credit.niso.org

TH: conceptualization, data curation, formal analysis, funding acquisition, methodology, project administration, resources, supervision, validation, visualization, writing – original draft

JB: conceptualization, investigation, methodology, writing – original draft

NR: conceptualization, investigation, methodology, writing – original draft

BH: conceptualization, supervision – original draft

XF: conceptualization, data curation, formal analysis, investigation, methodology, project administration, supervision, validation, visualization, writing – original draft

