## Supplementary Figures 1-12 for "Limb assignment, not spatial remapping, underlies tactile crossing effects"

for

#### Experiment 1

Supplementary Fig. 1 shows RT results of Experiment 1.
Supplementary Fig. 2 shows accuracy results of Experiment 1.
Table 1 shows Anova results of Experiment 1 including the factor SOA.

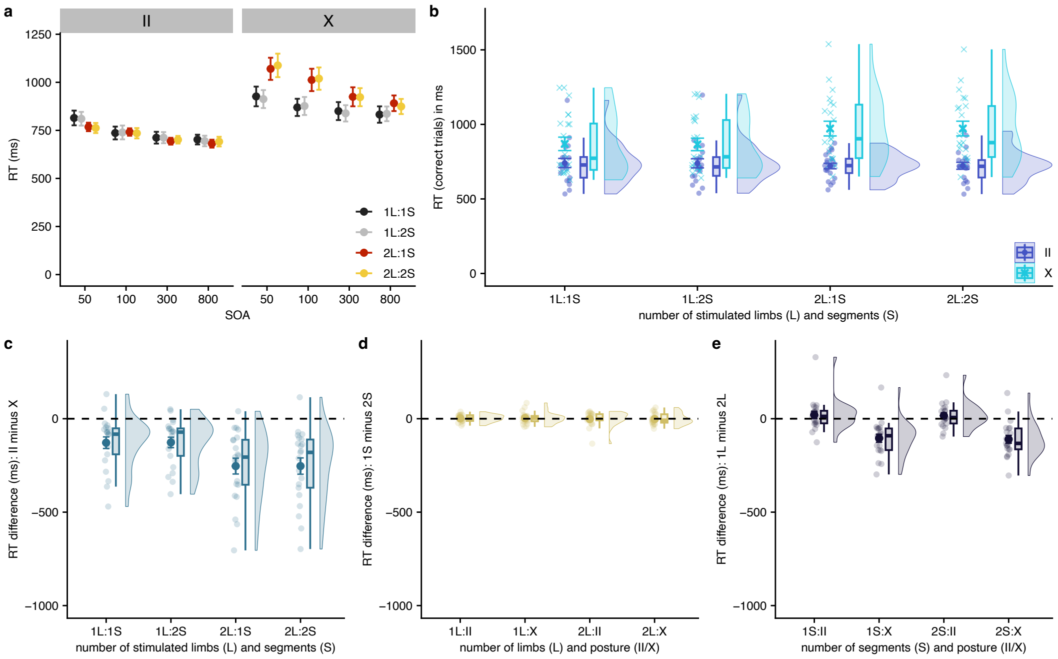


**Supplementary Figure 1: Reaction Time results of Experiment 1**. (a) Group performance split by all experimental factors: Limb Crossing (II: uncrossed, X: crossed); Number of Limbs (1 vs. 2 stimulated limbs); Number of Segments (1 vs. 2 limb segments stimulated); SOA, stimulus asynchrony in ms. (b) Uncrossed (dark blue) vs. crossed (light blue) performance across factors Number of Limbs and Number of Segments. (c) Difference scores of uncrossed minus crossed performance. (d) Difference scores of performance when stimuli occurred on a common segment, that is, the forearm, vs. on different segments, that is, fore- and upper arm. (e) Difference scores of performance when stimuli both occurred on one limb vs. when they occurred on two different limbs. The figure is equivalent to that of the main paper's Fig. 2 (BIS results of Experiment 1), and all conventions of that figure apply.

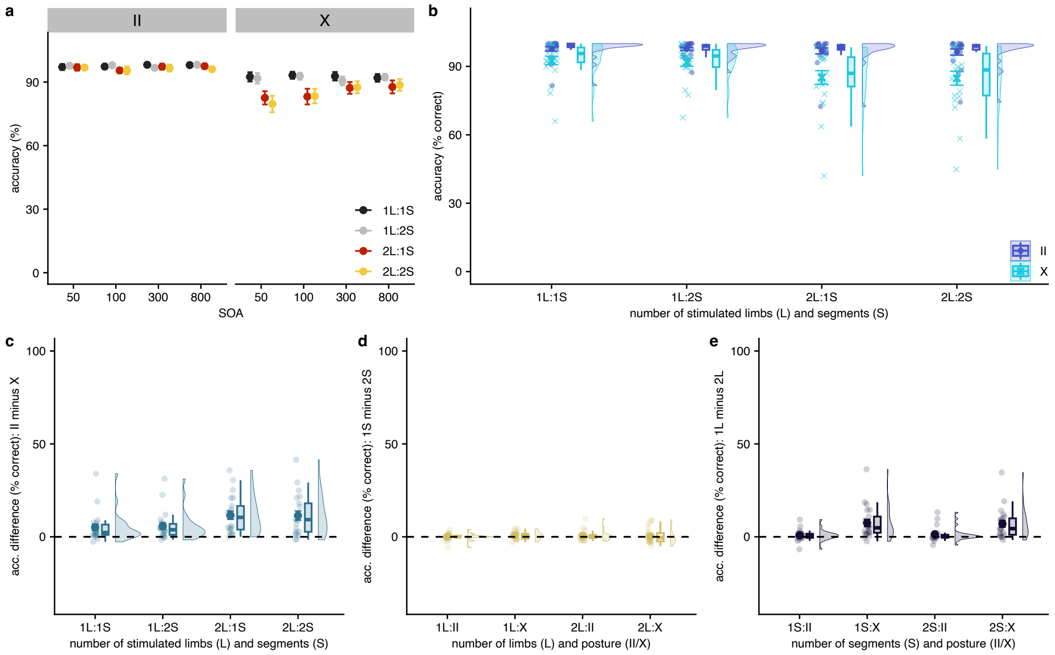


**Supplementary Figure 2: Accuracy results of Experiment 1**. (a) Group performance split by all experimental factors (see caption of Supplementary Figure 1). (b) Uncrossed (dark blue) vs. crossed (light blue) performance across factors Number of Limbs and Number of Segments. (c) Difference scores of uncrossed minus crossed performance. (d) Difference scores of performance when stimuli occurred on a common segment, that is, the forearm, vs. on different segments, that is, fore- and upper arm. (e) Difference scores of performance when stimuli both occurred on one limb vs. when they occurred on two different limbs. The figure is equivalent to that of the main paper’s Fig. 2 (BIS results of Experiment 1), and all conventions of that figure apply.

**SupplementaryTable 1: Results of the anova of Experiment 1 with factor SOA**. SOA had 4 levels (50, 100, 300, 800 ms). Degrees of freedom (df) of the SOA factor were corrected with the Greenhouse-Geisser method. The table lists corrected values.

| effect | df | F | $\eta^2$ | p |
|---|---|---|---|---|
| Posture | 1, 20 | 93.91 | 0.541 | <0.001 |
| Limbs | 1, 20 | 50.69 | 0.170 | <0.001 |
| Segments | 1, 20 | 0.63 | 0.001 | 0.436 |
| SOA | 1.66, 33.11 | 23.61 | 0.142 | <0.001 |
| Posture x Limbs | 1, 20 | 87.20 | 0.153 | <0.001 |
| Posture x Segments | 1, 20 | 0.02 | <0.001 | 0.884 |
| Limbs x Segments | 1, 20 | 0.18 | <0.001 | 0.675 |
| Posture x SOA | 2.65, 53 | 3.77 | 0.012 | 0.020 |
| Limbs x SOA | 2.38, 47.5 | 7.90 | 0.033 | 0.001 |
| Segments x SOA | 2.17, 43.48 | 0.57 | 0.001 | 0.584 |
| Posture x Limbs x Segments | 1, 20 | 2.55 | 0.002 | 0.126 |
| Posture x Limbs x SOA | 2.55, 51.07 | 10.08 | 0.036 | <0.001 |
| Posture x Segments x SOA | 2.18, 43.6 | 0.31 | 0.001 | 0.753 |
| Limbs x Segments x SOA | 2.48, 49.64 | 1.06 | 0.002 | 0.366 |
| Posture x Limbs x Segments x SOA | 2.63, 52.67 | 1.15 | 0.002 | 0.334 |

### Experiment 2

Supplementary Fig. 3 shows RT results of Experiment 2.

Supplementary Fig. 4 shows accuracy results of Experiment 2.

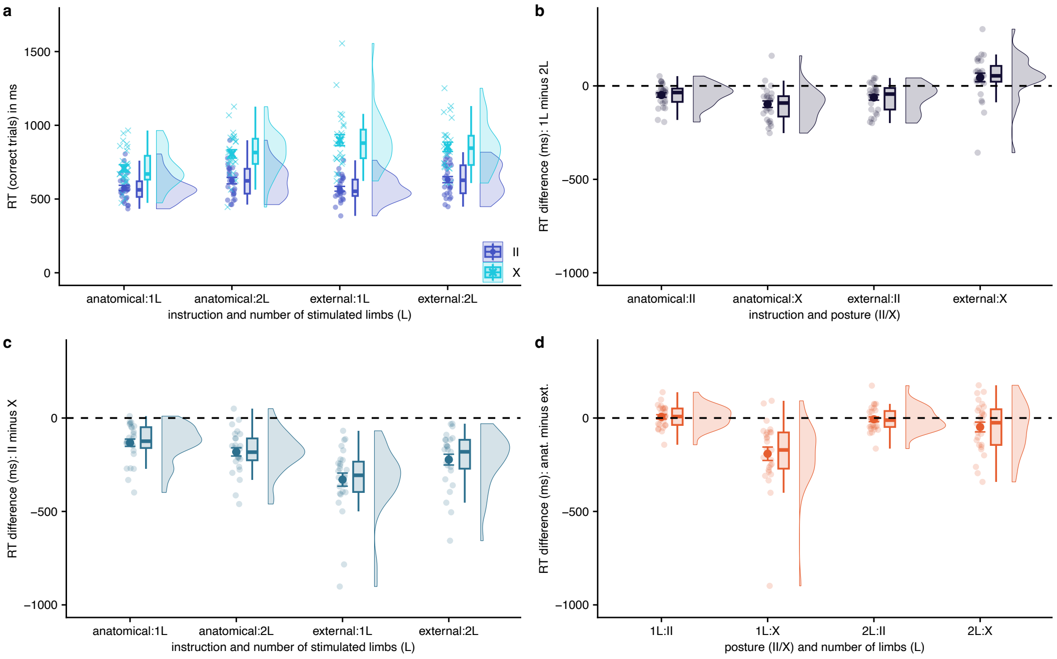


**Supplementary Figure 3: Reaction Time results of Experiment 2**. (a) Uncrossed (dark blue) vs. crossed (light blue) performance across factors Number of Limbs and Instruction. (b) Difference scores of performance when stimuli both occurred on one limb vs. when they occurred on two different limbs. (c) Difference scores of uncrossed minus crossed performance. (d) Difference scores of performance under anatomical vs. external instructions. The figure is equivalent to that of the main paper’s Fig. 4 (BIS results of Experiment 2), and all conventions of that figure apply.

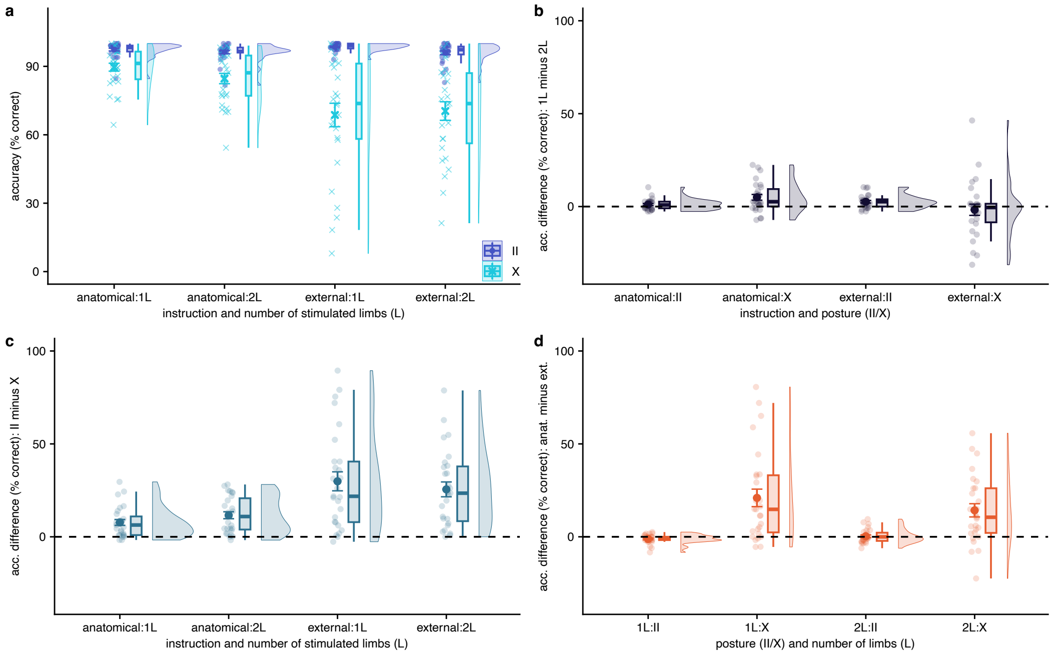


**Supplementary Figure 4: Accuracy results of Experiment 2**. (a) Uncrossed (dark blue) vs. crossed (light blue) performance across factors Number of Limbs and Instruction. (b) Difference scores of performance when stimuli both occurred on one limb vs. when they occurred on two different limbs. (c) Difference scores of uncrossed minus crossed performance. (d) Difference scores of performance under anatomical vs. external instructions. The figure is equivalent to that of the main paper’s Fig. 4 (BIS results of Experiment 2), and all conventions of that figure apply.

### Experiment 3

Supplementary Fig. 5 shows RT results of Experiment 3 (pooled across 3a and 3b).
Supplementary Fig. 6 shows accuracy results of Experiment 3 (pooled across 3a and 3b).
Supplementary Fig. 7 shows BIS results of Experiment 3a.
Supplementary Fig. 8 shows BIS results of Experiment 3b.

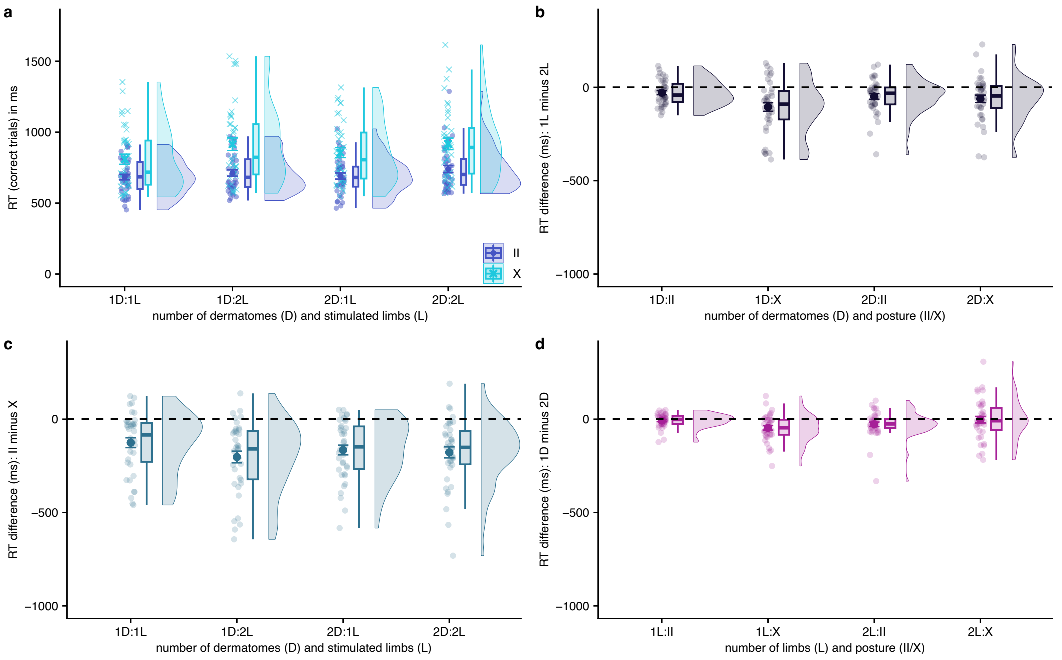


**Supplementary Figure 5: Reaction Time results of Experiment 3**. (a) Uncrossed (dark blue) vs. crossed (light blue) performance across factors Number of Limbs and Number of Dermatomes. (b) Difference scores of performance when stimuli both occurred on one limb vs. when they occurred on two different limbs. (c) Difference scores of uncrossed minus crossed performance. (d) Difference scores of performance for stimulus pairs in 1 vs. in 2 dermatomes. The figure is equivalent to that of the main paper’s Fig. 6 (BIS results of Experiment 3), and all conventions of that figure apply.

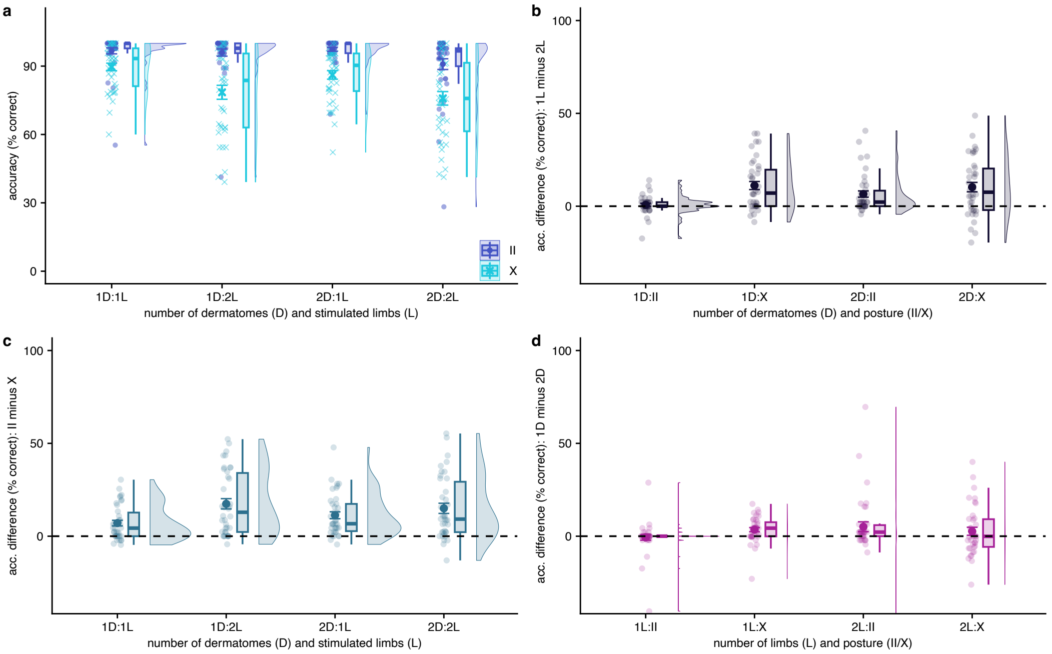


**Supplementary Figure 6: Accuracy results of Experiment 3**. (a) Uncrossed (dark blue) vs. crossed (light blue) performance across factors Number of Limbs and Number of Dermatomes. (b) Difference scores of performance when stimuli both occurred on one limb vs. when they occurred on two different limbs. (c) Difference scores of uncrossed minus crossed performance. (d) Difference scores of performance for stimulus pairs in 1 vs. in 2 dermatomes. The figure is equivalent to that of the main paper’s Fig. 6 (BIS results of Experiment 3), and all conventions of that figure apply.

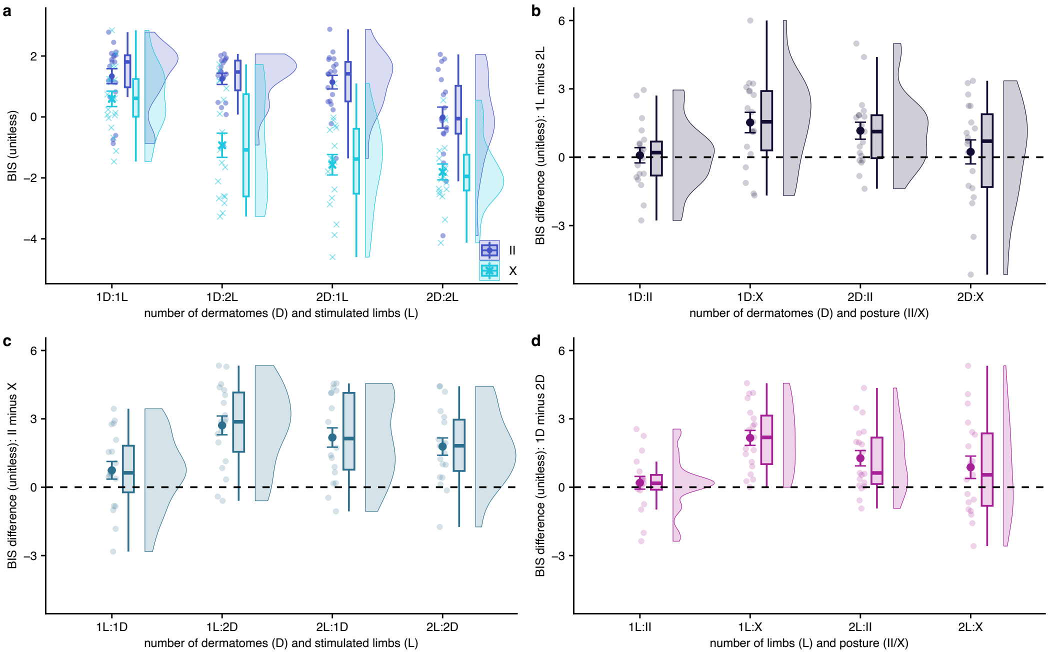


**Supplementary Figure 7: BIS results of Experiment 3a**. (a) Uncrossed (dark blue) vs. crossed (light blue) performance across factors Number of Limbs and Number of Dermatomes. (b) Difference scores of performance when stimuli both occurred on one limb vs. when they occurred on two different limbs. (c) Difference scores of uncrossed minus crossed performance. (d) Difference scores of performance for stimulus pairs in 1 vs. in 2 dermatomes. The figure is equivalent to that of the main paper’s Fig. 6 (BIS results of Experiment 3), and all conventions of that figure apply; however, we acquired two groups of participants with different definitions of 1 vs. 2 dermatome stimulus locations, which are reported pooled in the paper. See main paper’s Methods for details of the way dermatome locations were determined in Exp. 3a.

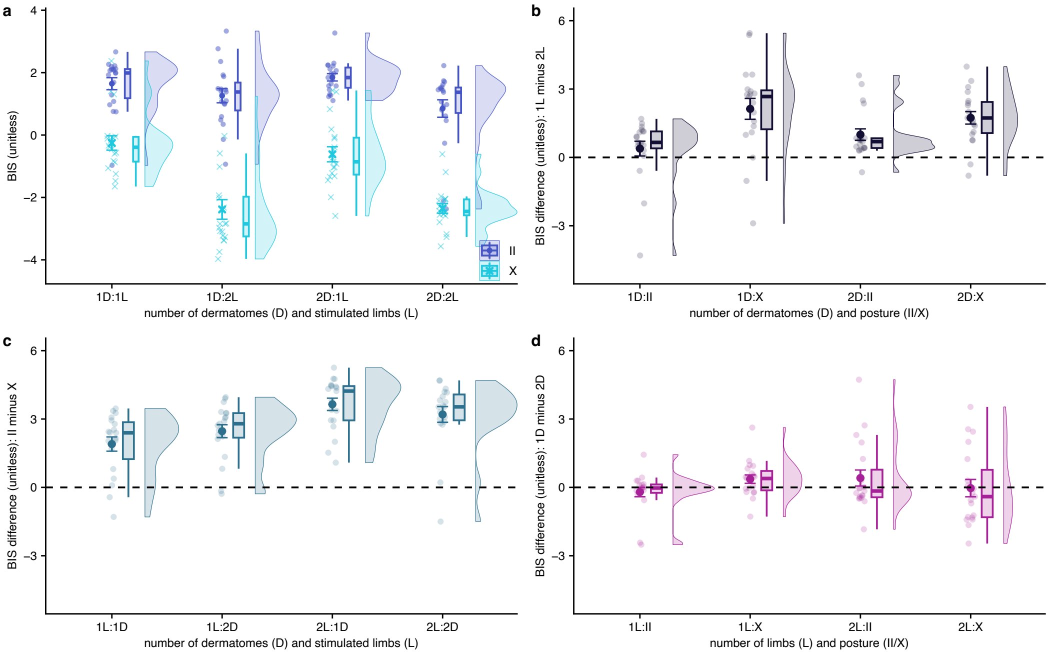


**Supplementary Figure 8: BIS of Experiment 3b**. (a) Uncrossed (dark blue) vs. crossed (light blue) performance across factors Number of Limbs and Number of Dermatomes. (b) Difference scores of performance when stimuli both occurred on one limb vs. when they occurred on two different limbs. (c) Difference scores of uncrossed minus crossed performance. (d) Difference scores of performance for stimulus pairs in 1 vs. in 2 dermatomes. The figure is equivalent to that of the main paper’s Fig. 6 (BIS results of Experiment 3), and all conventions of that figure apply; however, we acquired two groups of participants with different definitions of 1 vs. 2 dermatome stimulus locations, which are reported pooled in the paper. See main paper’s Methods for details of the way dermatome locations were determined in Exp. 3b.

### Experiment 4

Supplementary Fig. 9 shows RT results of Experiment 4.

Supplementary Fig. 10 shows accuracy results of Experiment 4.

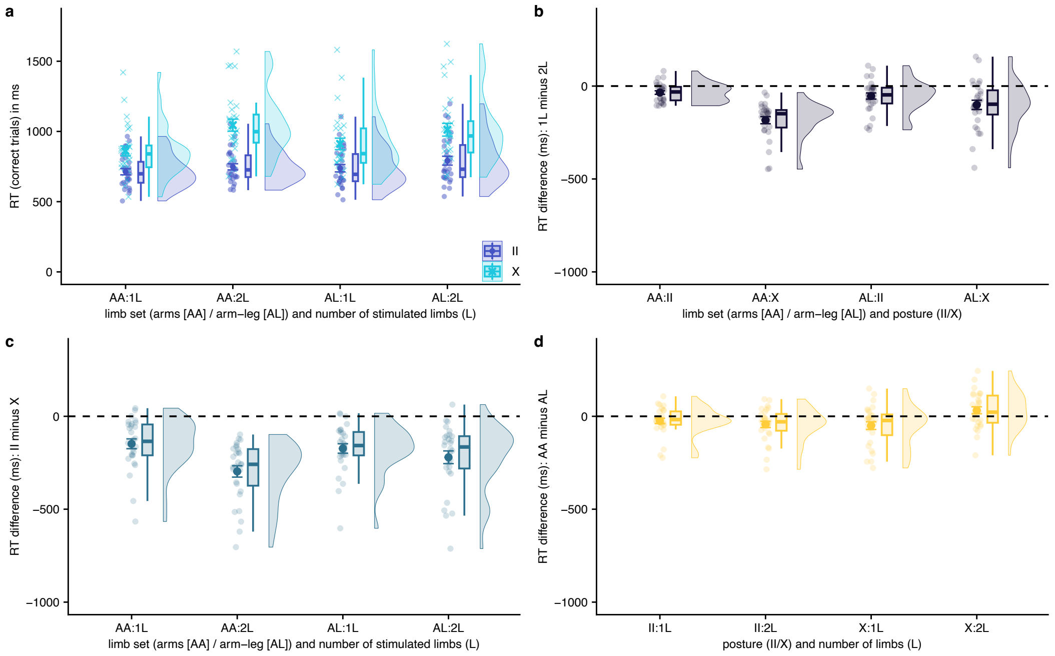


**Supplementary Figure 9: Reaction time results of Experiment 4**. (a) Uncrossed (dark blue) vs. crossed (light blue) performance across factors Limb Set and Number of Limbs. (b) Difference scores of performance when stimuli both occurred on one limb vs. when they occurred on two different limbs. (c) Difference scores of uncrossed minus crossed performance. (d) Difference scores of performance for the arm-arm vs. the arm-leg limb set. The figure is equivalent to that of the main paper’s Fig. 6 (BIS results of Experiment 3), and all conventions of that figure apply.

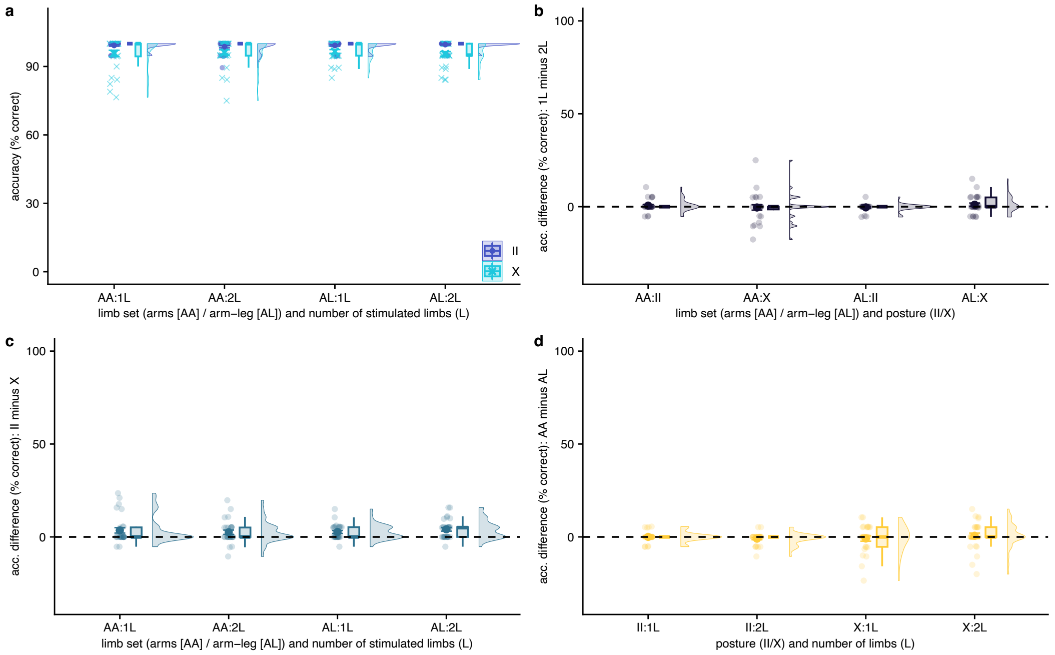


**Supplementary Figure 9: Accuracy results of Experiment 4**. (a) Uncrossed (dark blue) vs. crossed (light blue) performance across factors Limb Set and Number of Limbs. (b) Difference scores of performance when stimuli both occurred on one limb vs. when they occurred on two different limbs. (c) Difference scores of uncrossed minus crossed performance. (d) Difference scores of performance for the arm-arm vs. the arm-leg limb set. The figure is equivalent to that of the main paper’s Fig. 6 (BIS results of Experiment 3), and all conventions of that figure apply.

#### Experiment 5

We did not analyze RT in Experiment 5, because participants had to give two responses (pointing to each of the two stimuli). Thus, neither RT nor BIS measures exist.
